# Hierarchical divergence across genomic, phenotypic, and microbiome dimensions in two annual killifish species from Malawi

**DOI:** 10.64898/2026.07.26.740841

**Authors:** Tamanna Rahman Joya, Peyton K. Warren, W. Thompson Andrew, Enoch Ng’oma

**Affiliations:** Department of Biology, University of Texas at Arlington, TX 76019; Division of Biological Sciences, University of Missouri, Columbia, MO 65211; Department of Biological Sciences, Western Michigan University, Kalamazoo, MI 49008

**Keywords:** phenotypic divergence, regulatory evolution, microbiome, hydrological structure, multivariate morphology, seasonal killifish

## Abstract

Divergence among populations and species commonly occurs across multiple biological levels, yet the extent to which genomic, phenotypic, and ecological dimensions are coupled remains poorly understood. We integrated whole-genome sequencing, multivariate morphology, population-level phylogenetic structure, and gut microbiome composition to evaluate divergence in two allopatric annual killifish species across spatially structured populations in Malawi. Geographic and hydrological structure emerged as the primary axis of divergence, with strong differentiation between species and among populations. Genome-wide analyses revealed consistent clustering among populations in ordination and phylogenetic analyses, indicating pronounced spatial structuring of genomic variation. Although genomic divergence was widespread, exon-level enrichment analyses revealed distinct signatures across evolutionary scales. Divergence within *N. kirki* was associated primarily with translation-related functions, whereas divergence within *N. wattersi* involved ATP biosynthesis and physiological homeostasis. Interspecific divergence was enriched for transcription factor activity and transcription factor binding, implicating regulatory evolution as a major component of species differentiation. Morphological variation was likewise strongly structured among populations and aligned with drainage systems and geographic regions but was not correlated with genome-wide genetic differentiation, indicating partial decoupling between genotype and multivariate phenotype. Gut microbiome composition represented a more environmentally responsive layer of divergence, broadly reflecting host species and drainage structure while exhibiting greater overlap among populations. These results support a hierarchical model in which geographic and hydrological structure organize stable genomic divergence, whereas phenotypic and microbiome variation represent increasingly context-dependent and only partially aligned biological layers. Our findings highlight the value of integrating multiple biological levels to understand how evolutionary processes shape divergence in natural populations.

## Introduction

Divergence among populations and species commonly occurs across multiple biological levels, from genomic differentiation to phenotypic variation and host-associated biotic communities such as microbiomes. Genomic data can reveal signatures of historical isolation, heterogeneous differentiation, and locus-specific selection across the genome (Ellegren & Galtier, 2016; Nosil et al., 2009), while phenotypic data capture the integrated outcomes of development, ecology, and selection (Arnold, 1983; Lande, 1979). Microbiomes, in turn, represent a dynamic ecological interface between organisms and their environments, reflecting host and symbiont-associated processes and local environmental conditions (Ley et al., 2008; Shapira, 2016). Recent conceptual frameworks, such as the genotype-to-environment (G2E) perspective (Li et al., 2025), emphasize that these layers are linked through bidirectional interactions in which genomic variation shapes organismal and ecological states, while environmental feedbacks, including influence of microbial communities, modulate phenotypic expression and evolutionary trajectories. Understanding how these layers covary, or decouple, remains a central challenge in evolutionary ecology (Hendry, 2013; Stuart et al., 2017).

Linking genomic and phenotypic divergence remains one of the central challenges in evolutionary biology because phenotypic evolution is inherently multivariate and arises through interactions among genetic, developmental, and environmental processes. Complex traits are typically influenced by numerous loci distributed across the genome, producing coordinated changes in suites of traits rather than simple one-trait/one-locus relationships (Boyle et al., 2017; Rockman, 2012). Increasing evidence further indicates that phenotypic evolution frequently reflects changes in gene regulation, including *cis*- or *trans*-regulatory mechanisms that alter the spatial or temporal expression of genes rather than modifications to downstream protein-coding sequences (Carroll, 2008; King & Wilson, 1975; Wray, 2007). Consequently, phenotypic divergence often exhibits only partial or non-linear correspondence with genome-wide differentiation because it is additionally shaped by ecological context, developmental plasticity, and sex-specific processes (Leinonen et al., 2013; Pfennig et al., 2010; West-Eberhard, 2003). These interactions imply that phenotypes emerge from the combined effects of genomic variation and heterogeneous environmental inputs rather than direct mappings between genotype and trait values. Consequently, the biological pathways underlying divergence need not be conserved across evolutionary scales. Population- and species-level differentiation may instead involve distinct physiological, metabolic, developmental, or regulatory processes (Signor & Nuzhdin, 2018; Stern & Orgogozo, 2008). Identifying functional classes of genes that contribute disproportionately to divergence therefore provides an important bridge between genome-wide patterns of differentiation and their potential phenotypic consequences.

Geographic and environmental heterogeneity can structure divergence across multiple biological levels, but these relationships often depend on evolutionary scale. Population differentiation may closely reflect hydrological structure, historical connectivity, and barriers to gene flow (Avise, 2000; Hughes et al., 2009), even when morphological differentiation is only weakly associated with geographic distance or a limited set of measured environmental variables (Marroig & Cheverud, 2005). Similarly, host-associated microbiomes may retain signatures of host population structure (Brooks et al., 2016) while responding rapidly to local environmental conditions (Rothschild et al., 2018; Sullam et al., 2012), producing patterns that are only partially aligned with host genomic and phenotypic variation (Araujo et al., 2026; Henry et al., 2021). In hydrologically structured allopatric systems, where geographic isolation and restricted connectivity are the dominant drivers of divergence, genomic, phenotypic, and microbiome structure are therefore expected to exhibit broadly similar spatial patterning (Avise, 2000; Manel et al., 2003; Moeller et al., 2016). However, because phenotypes and microbiomes also respond to local ecological conditions, developmental plasticity, and host physiology, these biological layers need not covary perfectly with genome-wide differentiation. Such scale-dependent patterns of alignment and decoupling are consistent with G2E expectations that different biological levels integrate genomic and environmental signals over distinct spatial and temporal contexts (Li et al., 2025).

Killifishes of the genus *Nothobranchius* are annual species distributed across eastern Africa, occupying ephemeral pools that form during the rainy season. These habitats are embedded within hydrologically structured landscapes and impose strong seasonal bottlenecks, rapid life histories, and repeated local isolation, all of which promote population divergence (Genade et al., 2005; Terzibasi et al., 2008; Valdesalici & Cellerino, 2003). Consistent with these ecological features, genomic studies in *Nothobranchius* have revealed substantial population structure and signatures of adaptation across environmental gradients, including differentiation associated with aridity and life-history traits (Bartáková et al., 2013; Cui et al., 2019). This system provides a unique opportunity for evaluating how repeated isolation, environmental heterogeneity, and rapid life histories contribute to population divergence. The availability of genome-wide data further allows assessment of whether divergence is concentrated in particular genomic regions and biological functions or distributed broadly across the genome providing insight into the genomic architecture of divergence in naturally fragmented populations.

In this study, we focus on two allopatric species of annual killifish in Malawi distributed in two hydrologically distinct regions: the Chilwa-Chiuta depression and the Lake Malawi-upper Shire River basin. Integrating molecular and morphological data, Ng’oma et al. (2013) demonstrated that these basin-associated populations represent distinct evolutionary lineages, formally recognizing the Lake Malawi-upper Shire populations as *N. wattersi*, separate from *N. kirki* of the Chilwa-Chiuta system. This divergence is consistent with the geomorphological history of basin isolation associated with the development of the East African Rift system (Delvaux, 1995; Lancaster, 1981; Lister, 1965; Shroder, 1972). In contrast, the Chilwa and Chiuta systems remained connected until relatively recently (∼ 9 - 6 ka) and may have experienced intermittent hydrological connectivity into historical times (Lancaster, 1981; Nicholson, 1998; Thomas et al., 2009), generating expectations of comparatively shallow or incomplete divergence between populations in these basins. At finer spatial scales, topographic and hydrological heterogeneity may further structure divergence within basins. In particular, upland populations in the Namwera Hills north of Lake Chiuta occupy a distinct drainage and elevational setting that may restrict connectivity with surrounding floodplain habitats, potentially promoting additional population differentiation within *N. kirki*. These hydrologically structured landscapes provide a natural framework for testing how historical basin isolation, contemporary drainage structure, and local environmental variation shape divergence across genomic, phenotypic, and ecological dimensions.

Here, we investigate how previously identified divergence (Ng’oma et al., 2013) is structured across biological levels in a spatially heterogeneous system, and whether these layers show consistent or partially decoupled spatial patterning. To do so, we integrate whole-genome sequencing, multivariate analyses of morphology, population-level phylogenetic structure, and gut microbiome data across matched populations of *N. kirki* and *N. wattersi*. We aim to (i) characterize spatial structuring of divergence across the G2E axis, (ii) evaluate the extent to which genomic, phenotypic, and microbiome divergence are aligned across populations, and (iii) assess whether divergence at population and species levels is associated with distinct functional signatures in protein-coding genes and regulatory pathways. Specifically, we test whether divergence is similarly structured across biological levels or whether different layers reflect distinct ecological and evolutionary processes operating over different spatial and temporal contexts. We hypothesized that genomic divergence would most strongly reflect hydrological and geographic isolation, whereas phenotypic and microbiome variation would show greater sensitivity to local ecological and environmental conditions. We further predicted that functional enrichment analyses would reveal scale-dependent signatures of divergence, with distinct biological pathways contributing to differentiation within and between species.

## Materials and Methods

### Study system and sampling

We sampled populations of two allopatric species of Nothobranchius across their geographic ranges in Malawi (Fig. 1). *Nothobranchius kirki* Jubb inhabits the recently isolated Chilwa-Chiuta drainage in southeastern Malawi, whereas *N. wattersi* (Ng’oma et al., 2013) occurs in the southern Lake Malawi and upper Shire River floodplains. Sampling was designed to capture major hydrological and geographic discontinuities expected to structure population divergence. Biological datasets comprised whole-genome sequencing, morphology, and gut microbiome data collected from populations sampled within the same localities or individual pools whenever possible (Table 1). Although coverage differed somewhat among datasets, the shared population-based sampling framework enabled comparisons of divergence across biological levels while acknowledging that incomplete overlap limited direct matrix-based comparisons for some analyses.

**Figure 1:**
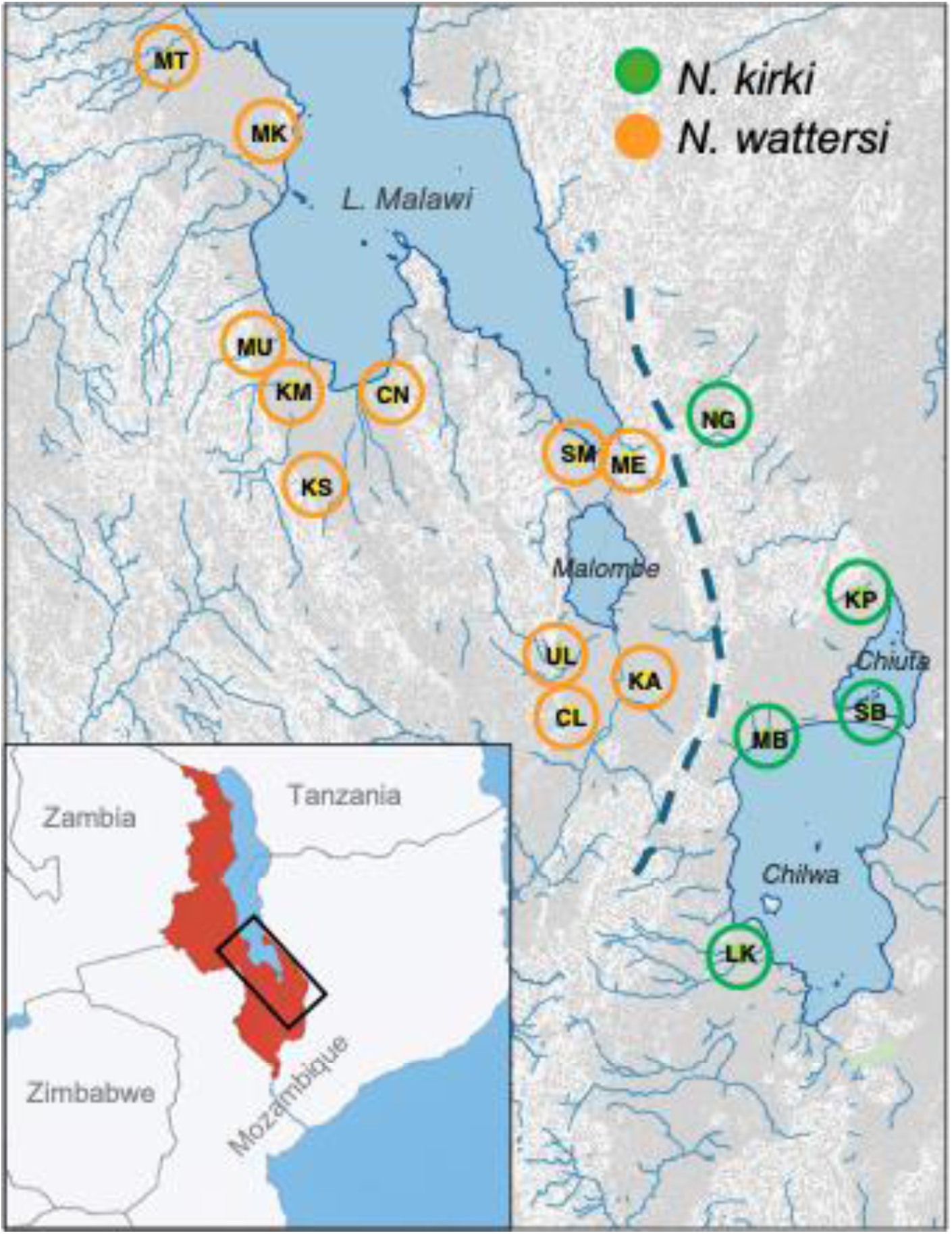
Range-wide sampling of *N. kirki* (green circles) from lakes Chilwa and Chiuta depressions and *N. wattersi* (orange circles) from L. Malawi and Shire River flood plains. We found no specimens of *N. wattersi* in the most northerly historical collection area (*N.* spec. “Benga” MW 88/12, Brian Watters, Ng’oma et al. (2013)) north of MT location despite significant sampling effort. Insert: regional context of the country Malawi (red, with Lake Malawi, blue), and the general distribution of species studied here (black ellipse).

**Table 1:**
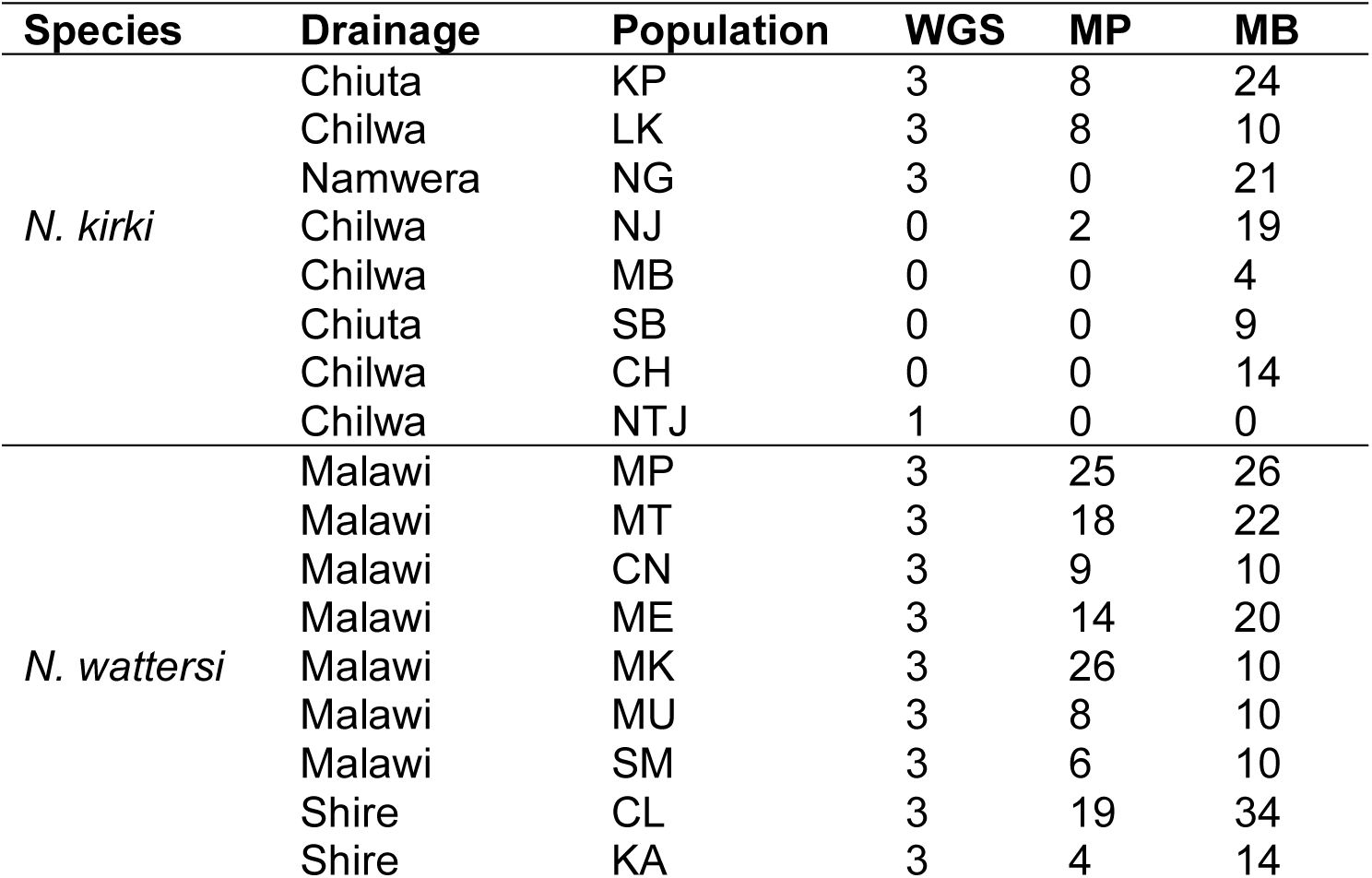
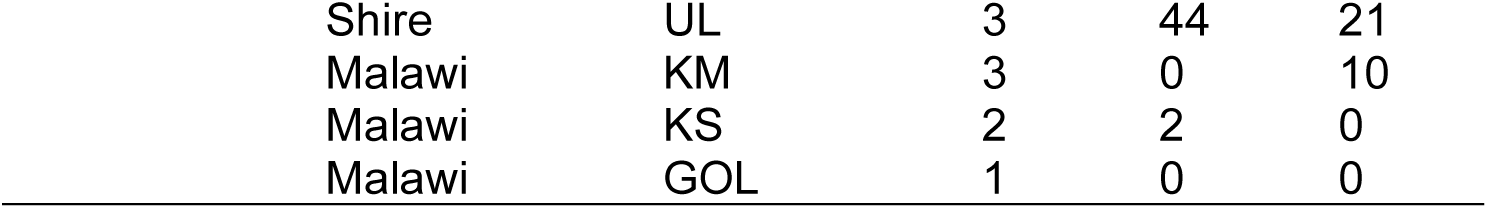
Sampling design and dataset coverage across populations of *N. kirki* and *N. wattersi*. Values indicate the number of individuals available for whole-genome sequencing (WGS), morphology (MP), and gut microbiome (MB) analyses. Integration among datasets was performed at the population (sampling pool) level rather than through matching individuals across datasets.

| Species | Drainage | Population | WGS | MP | MB |
| --- | --- | --- | --- | --- | --- |
| *N. kirki* | Chiuta | KP | 3 | 8 | 24 |
|  | Chilwa | LK | 3 | 8 | 10 |
|  | Namwera | NG | 3 | 0 | 21 |
|  | Chilwa | NJ | 0 | 2 | 19 |
|  | Chilwa | MB | 0 | 0 | 4 |
|  | Chiuta | SB | 0 | 0 | 9 |
|  | Chilwa | CH | 0 | 0 | 14 |
|  | Chilwa | NTJ | 1 | 0 | 0 |
| *N. wattersi* | Malawi | MP | 3 | 25 | 26 |
|  | Malawi | MT | 3 | 18 | 22 |
|  | Malawi | CN | 3 | 9 | 10 |
|  | Malawi | ME | 3 | 14 | 20 |
|  | Malawi | MK | 3 | 26 | 10 |
|  | Malawi | MU | 3 | 8 | 10 |
|  | Malawi | SM | 3 | 6 | 10 |
|  | Shire | CL | 3 | 19 | 34 |
|  | Shire | KA | 3 | 4 | 14 |

|  |  |  |  |  |
| --- | --- | --- | --- | --- |
| Shire | UL | 3 | 44 | 21 |
| Malawi | KM | 3 | 0 | 10 |
| Malawi | KS | 2 | 2 | 0 |
| Malawi | GOL | 1 | 0 | 0 |

### Whole genome sequence analysis

#### DNA isolation, library preparation, and sequencing

Total genomic DNA was extracted from fin tissue of ethanol-preserved specimens (*N. kirki*, n = 10; *N. wattersi*, n = 36) using the Zymo Quick-DNA™ Miniprep Plus Kit following the manufacturer’s protocol. DNA quantity and quality were assessed using NanoDrop spectrophotometry and agarose gel electrophoresis. Sequencing libraries were prepared with the Illumina DNA Prep kit using dual-index adapters and size-selected to an average insert size of approximately 550 bp. Libraries were quantified using a Qubit fluorometer and fragment size distributions verified using an Agilent Fragment Analyzer before equimolar pooling. Sequencing was performed at the University of Missouri Genomics Technology Core on an Illumina NovaSeq X platform (150-bp paired-end reads). Libraries were distributed across three sequencing lanes to achieve a target yield of approximately 45 Gb per sample, after which lane-level reads were merged prior to downstream analyses.

#### Bioinformatic processing and phylogenomic inference

Sequencing reads were processed using two complementary computational workflows. One generated a filtered SNP dataset for population genomic analyses, whereas the second used the raw sequencing reads for phylogenomic inference. Population genomic analyses were performed by the University of Missouri Bioinformatics and Analytics Core using a custom Nextflow workflow (Di Tommaso et al., 2017). Reads were aligned to the *Nothobranchius furzeri* reference genome (Reichwald et al., 2015) using Minimap2 (H. Li, 2018). Duplicate reads were marked with GATK MarkDuplicates (v4.2.6.1), and variants were called for each sample using GATK HaplotypeCaller (Poplin et al., 2018). SNPs and indels were filtered using GATK VariantFiltration following best-practice recommendations (DePristo et al., 2011). Read-mapping statistics for all samples are provided in Table S3. The resulting filtered variant call format (VCF) file served as the basis for all downstream population genomic analyses.

For phylogenomic inference, raw FASTQ files from the same sequencing libraries were processed independently. Reads were demultiplexed using bcl-convert v3.8.2 (Illumina Inc.), adapter trimmed and quality filtered with fastp v0.23.1 (Chen et al., 2018), and lane-level FASTQ files were merged for each individual. Overlapping paired-end reads were merged using BBMerge (Bushnell et al., 2017), after which merged and unmerged reads were combined into a single FASTQ file per sample. Phylogenetic inference was performed using the WASTER pipeline (Zhang & Nielsen, 2026), with each individual treated as an independent operational taxonomic unit to avoid assumptions of species-level monophyly. Reads from single individuals representing 11 additional aplocheiloid species were retrieved from the NCBI Sequence Read Archive (ERX541553; SRR8877970, SRR8881303, SRR8881332, SRR8881054, SRR8881305, SRR8881337, SRR8881061, SRR8881307, SRR8881339, SRR8881063, and SRR8881310), processed identically, and included as outgroups for phylogenetic inference.

#### Population genomic analyses

Population genomic analyses were performed from the filtered whole-genome SNP dataset. To generate a high-quality dataset while retaining informative polymorphism, variants were filtered to retain sites with a minor allele count (MAC) ≥ 2 and site-level missingness ≤ 0.20, yielding a dataset with a mean site missingness of approximately 0.007. Sample identifiers were reconciled with associated metadata, the variant call format (VCF) file was converted to PLINK format, and sex metadata were incorporated into the binary files. To minimize the influence of linkage disequilibrium on multivariate analyses, SNPs were pruned using a 50-SNP sliding window, a 5-SNP step size, and an r^2^ threshold of 0.2, retaining approximately 4.2 million SNPs for downstream analyses.

Principal component analysis (PCA) was used to characterize genomic structure across the complete dataset and repeated separately within each species to resolve finer population structure. Genetic differentiation was quantified using F_ST_-based approaches. Species-level divergence was summarized using the Weir and Cockerham estimator (Weir & Cockerham, 1984), whereas within-species population structure was evaluated using pairwise population F_ST_ and genome-wide summary statistics. Genome-wide window-based scans were used to identify highly differentiated genomic regions, with the upper 2% of the F_ST_ distribution defining candidate regions for downstream analyses.

To characterize coding sequence divergence, SNPs overlapping annotated exons in the *N. furzeri* reference genome (Reichwald et al., 2015) were assigned to genes based on exon coordinates. Gene level divergence was summarized as the mean F_ST_ across all exonic SNPs associated with each gene and calculated separately for interspecific (*N. kirki* vs. *N. wattersi*) and within-species population comparisons. Genes represented by fewer than three exonic SNPs were excluded to reduce the influence of sparsely sampled loci. The resulting gene level divergence estimates were used for downstream functional enrichment analyses and visualization of genomic patterns of differentiation.

#### Functional enrichment analyses

Functional enrichment analyses were conducted using Gene Ontology (GO) annotations derived from the *N. furzeri* reference genome. We evaluated functional patterns using two complementary approaches. First, genes overlapping highly differentiated genomic windows identified from genome-wide F_ST_ scans were analyzed using over-representation analysis (ORA) implemented in clusterProfiler (Xu et al., 2024). Enrichment was evaluated separately for Biological Process (BP) and Molecular Function (MF) ontologies, with statistical significance assessed using the Benjamini-Hochberg false discovery rate (FDR) correction.

To evaluate coding-sequence divergence at the gene level, gene set enrichment analysis (GSEA) was performed using genes ranked by centered mean exon F_ST_ values generated from the population genomic analyses described above. Analyses were conducted separately for BP and MF ontologies using clusterProfiler, with significance assessed using Benjamini-Hochberg FDR correction. For significantly enriched gene sets, leading-edge genes were extracted to identify loci contributing most strongly to enrichment signals and were subsequently used to visualize genomic distributions and compare divergence levels relative to the genomic background.

### Morphological phenotype analysis

#### Morphometric measurements

Morphological variation was quantified from 18 linear morphometric traits (Table S1) measured with digital calipers (General® UltraTech®) on 193 individuals representing three populations of *N. kirki* (n = 18) and 11 populations of *N. wattersi* (n = 175; Table S2). Trait measurements were log_10_-transformed prior to analysis to improve normality and linearize allometric relationships. Measurement repeatability was assessed using fresh and ethanol-preserved specimens of laboratory-maintained *N. melanospilus* together with wild *N. kirki* and *N. wattersi*. Because several populations were sampled repeatedly within a season, each sampling event was assigned a unique identifier and incorporated into subsequent size-correction analyses.

#### Size correction and allometric scaling

Because our primary objective was to compare body shape independent of overall body size, we first evaluated whether allometric scaling relationships differed among species and populations. Length–mass allometry was examined using log-transformed total length (*TL*) and wet mass (*W*), restricting analyses to populations with *n* ≥ 5 to ensure reliable estimation of population-specific scaling relationships. We first compared nested models testing species-level differences:

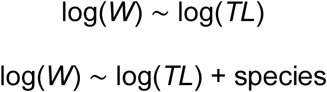

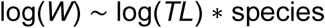

Analogous models were then used to test population-level heterogeneity in allometric scaling:

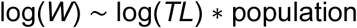

Model comparisons were conducted using ANOVA F-tests. Significant interaction terms for both species and population indicated heterogeneous allometric slopes among groups, making a single global size correction inappropriate.

Size correction was therefore performed separately for each species using population-specific linear models that included total length (*TL*), visit day, and sampling event as predictors of body mass. Predicted body mass from these models was used as a population-specific size factor, and morphometric traits were residualized against this factor to generate size-independent measures of body shape for downstream analyses. To evaluate potential measurement bias, we tested for observer effects using PERMANOVA with population and observer identity as predictors.

Population accounted for most of the multivariate variation (R^2^ = 0.62, *p* < 0.001), whereas observer identity explained less than 0.5% of the variance and was not significant (*P* = 0.14). Observer identity was therefore excluded from subsequent analyses.

### Geographic and environmental drivers of morphological divergence

#### Multivariate morphological variation

Patterns of morphological variation were characterized using principal components analysis (PCA) performed separately for each species on centered and scaled, size-corrected morphometric traits. Principal axes explaining at least 80% of the cumulative variance were retained for subsequent analyses. Trait loading coefficients were examined to identify the morphological variables contributing most strongly to each axis, and retained PC scores were used to quantify multivariate morphological divergence in all downstream analyses.

### Geographic and environmental structuring of morphology

We evaluated whether morphological divergence increased with geographic and environmental dissimilarity using Mantel tests conducted separately for each species. Morphological distances among populations were calculated as Euclidean distances in the retained morphological PCA space. Geographic distances were computed from field collected latitude and longitude coordinates using the Haversine great-circle formula (Earth radius = 6,371 km, (Sinnott, 1984)). Environmental variables (e.g., elevation, water temperature, pH) were centered and scaled, summarized using PCA, and Euclidean distances among populations were calculated from the retained environmental principal components. Mantel tests were then used to assess correlations between morphological and geographic distance matrices, and between morphological and environmental distance matrices, using Pearson correlations with 9,999 permutations. All analyses were performed in the vegan package (Oksanen et al., 2026) in R v4.6.0 (R Core Team, 2026).

#### Environmental predictors of morphological variation

To identify environmental variables associated with multivariate body shape, we conducted redundancy analysis (RDA) for *N. wattersi*, the only species with sufficient population level replication. Centered and scaled, size-corrected morphometric traits were used as response variables, while a subset of centered and scaled environmental variables (elevation, water temperature, and pH) served as explanatory variables.

Statistical significance was assessed using permutation tests (9,999 permutations), and variance inflation factors were examined to evaluate multicollinearity among predictors. RDA analyses were implemented using the rda() and anova.cca() functions in the vegan package.

#### Sexual dimorphism

Because the primary analyses focused on species- and population-level divergence, sexual dimorphism was evaluated separately for *N. wattersi*, the only species with sufficient and balanced sample sizes. Centered and scaled, size-corrected traits were used to calculate Euclidean distances among individuals, followed by principal coordinates analysis (PCoA). Group centroids and dispersion were estimated using betadisper() in vegan, and differences between sexes were evaluated using Welch’s t-tests on scores from the first PCoA axis, which captured most of the between-sex variation.

#### Sensitivity analysis

To account for potential non-independence among populations arising from shared evolutionary history (Felsenstein, 1985), we repeated morphology–environment analyses using partial redundancy analysis at the population level. Mean size-corrected morphometric traits for each population were analyzed with environmental principal components as explanatory variables and genomic principal components derived from whole-genome SNP data as conditioning variables. Variance partitioning was subsequently used to quantify the relative contributions of environmental and genomic structure to morphological variation.

### Microbiome analyses

#### 16S rRNA sequencing and read processing

For microbiome analyses, total DNA was extracted from whole-gut tissues using the Zymo Quick-DNA™ Miniprep Plus Kit (Cat. D4069; lots 239061 and 239505). Samples were homogenized using a Qiagen TissueLyser II with two 3.2 mm stainless steel beads (Biospec), incubated overnight, and processed according to the manufacturer’s protocol. DNA quality and concentration were assessed using a NanoDrop ND-1000 spectrophotometer and a Qubit 4 Fluorometer (Thermo Fisher Scientific), respectively.

Amplicon libraries targeting the V4 region of the bacterial 16S rRNA gene were sequenced on an Illumina MiSeq platform (paired-end 2 × 250 bp) using primers 515F and 806R.

Sequence processing was performed in R using the DADA2 pipeline (Callahan et al., 2016). Reads were quality filtered and trimmed to 210 bp, chimeric sequences were removed, and taxonomy was assigned using the SILVA v138.2 reference database. Of the 288 original samples, 238 passed filtering and sequence assembly. Processed reads were imported into a phyloseq object together with sample metadata. Host-derived and mitochondrial sequences were removed by excluding taxa outside Bacteria and Archaea, potential contaminants were identified using the frequency-based method implemented in the decontam R library, and amplicon sequence variants (ASVs) occurring in fewer than two samples were excluded. Sequence counts were converted to relative abundances for downstream analyses.

#### Microbiome community analyses

Microbial community composition was summarized using relative abundance profiles generated with phyloseq. Community structure was evaluated using principal coordinates analysis (PCoA) based on weighted Bray–Curtis dissimilarities. Ordinations were performed separately for each species to characterize population- and sex-associated variation in microbiome composition, with 75% confidence ellipses drawn around group centroids for visualization.

## Results

### Species-level divergence

#### Genomic divergence

Genomic analyses revealed strong separation between *N. kirki* and *N. wattersi*. PCA based on approximately 4.2 million linkage-disequilibrium-pruned SNPs clearly separated the two species, with PC1 and PC2 explaining 27.7% and 10.1% of genomic variation, respectively (Fig. 2a). The species-level PCA also revealed additional intraspecific structure, particularly within *N. kirki*, indicating that genome-wide SNP variation captures both interspecific and intraspecific differentiation (Fig. 2b,c).

**Figure 2:**
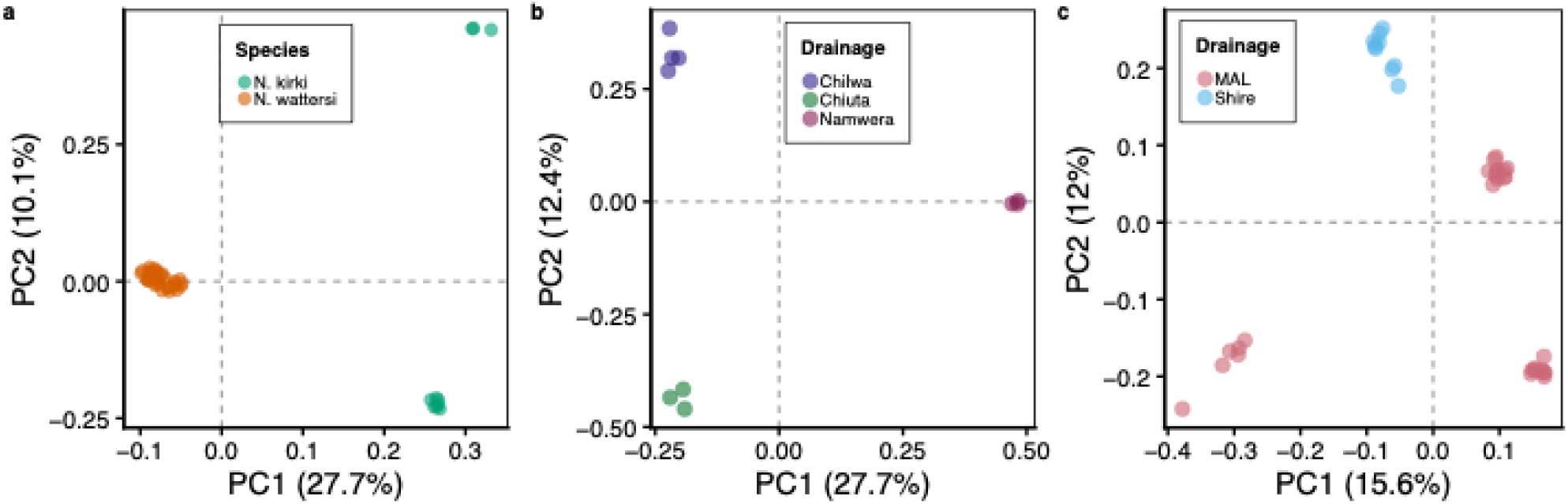
Principal components analysis (PCA) of genomic variation. **(a)** Inter-species differentiation between *Nothobranchius kirki* and *N. wattersi*. **(b, c)** Population structure within *N. kirki*, and *N. wattersi*, respectively. *N. kirki*, n = 10; *N. wattersi*, n = 36. MAL, Lake Malawi basin.

Mean genome-wide differentiation between species was moderate (mean F_ST_ = 0.221), but highly heterogeneous across loci (median = 0.117, 2.5-97.5%; range: −0.051 to 1.0). The weighted genome-wide F_ST_ estimate was substantially higher (0.432), indicating that highly differentiated loci contributed disproportionately to overall genomic differentiation.

Exon-level analyses revealed stronger functional organization than window-based scans. A total of 906 genes were highly differentiated (upper 2% of the F_ST_ distribution, Table S4). Over-representation analyses identified few GO categories that remained significant after multiple-testing correction, including thyroxine 5’-deiodinase activity (*p* <0.05, Table S5), suggesting that highly differentiated coding loci were distributed across numerous biological functions. In contrast, gene set enrichment analyses (GSEA) of ranked exon-level divergence scores recovered significant biological process (BP) and molecular function (MF) categories across all evolutionary scales examined (Fig. 3, Table 2, Table S6). Importantly, mean F_ST_ values were not significantly influenced by variation in gene size or SNP density, indicating that these factors did not bias GSEA results (Fig. S1). Within *N. kirki*, enriched categories were dominated by translation initiation, protein synthesis, ion transport and homeostasis. Within *N. wattersi*, ATP biosynthetic processes were positively enriched, whereas nucleosome assembly, oxygen transport, transcriptional initiation, and myelination were negatively enriched, indicating that these categories were associated with genes exhibiting comparatively low exon divergence. Between species, enrichment was concentrated in DNA binding and transcription factor binding (Fig. 3a), together with additional categories related to cellular organization and signaling. Collectively, these enriched categories point to coordinated divergence in pathways associated with energy metabolism, gene regulation, and physiological maintenance, while conservation of chromatin organization, oxygen transport, and neural maintenance suggests functional constraint on core processes underlying organismal homeostasis.

**Figure 3:**
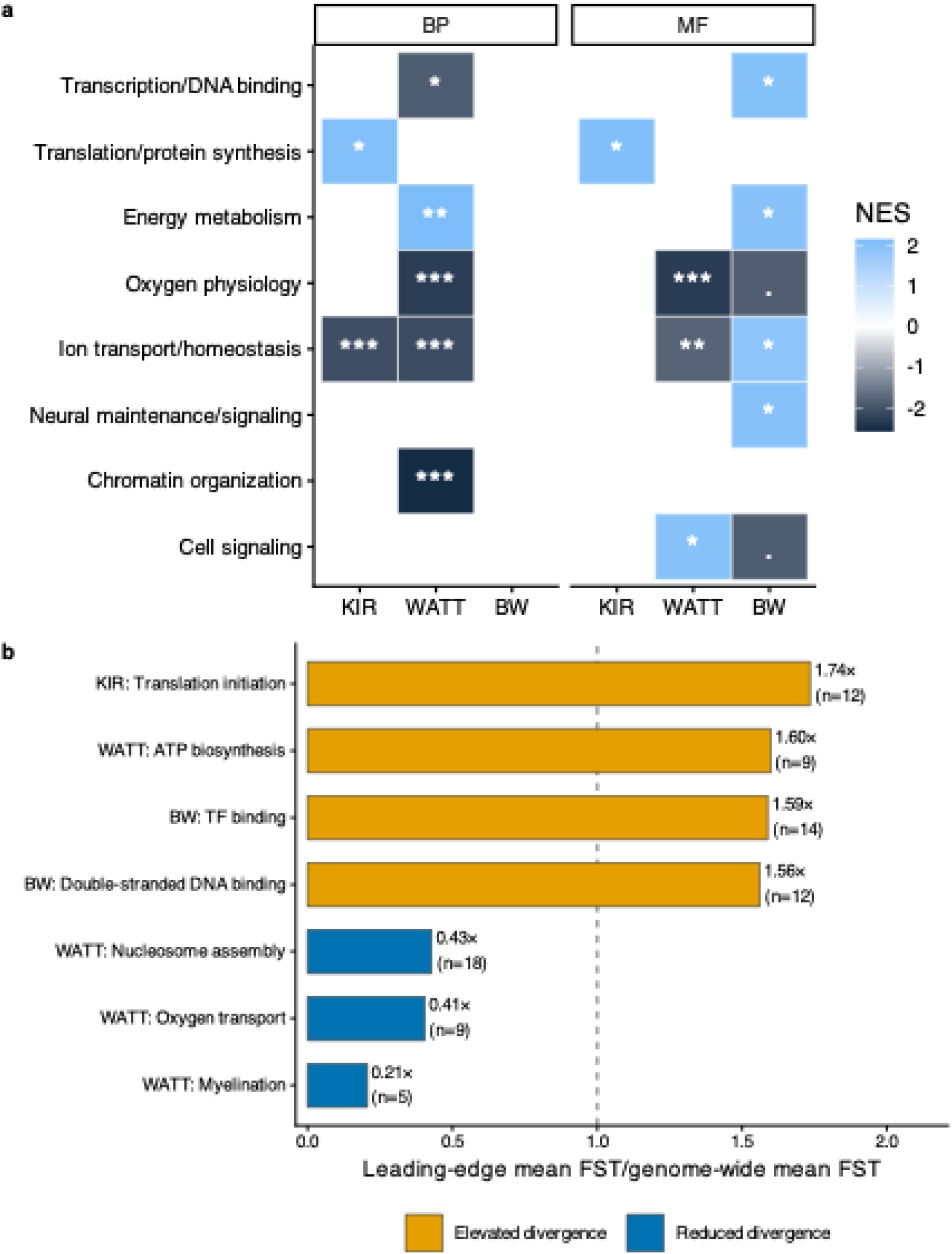
Functional themes associated with exon-level genomic divergence across hierarchical evolutionary scales. **(a)** Heatmap summarizing significantly enriched Gene Ontology (GO) biological process (BP) and molecular function (MF) categories identified by gene set enrichment analysis (GSEA) using *N. furzeri* annotations. Individual GO terms were grouped into broader biological themes. Cell color indicates the normalized enrichment score (NES). Symbols within cells indicate significance level of GO term contributing to each theme (FDR : *** < 0.001, ** < 0.01, * < 0.05, • < 0.10). **(b)** Relative exon-level divergence of leading-edge genes. Mean exon F_ST_ of leading-edge genes contributing to significant GSEA enrichments expressed relative to the genome-wide mean exon F_ST_ for each contrast. Bars indicate the ratio of mean exon F_ST_ in leading-edge genes to the corresponding genome-wide background. Values adjacent to bars indicate fold differences, and sample sizes (n) indicate the number of leading-edge genes contributing to each functional category.

**Table 2:** Principal functional themes associated with exon-level divergence. Representative GO categories identified by GSEA are shown for within-*N. kirki* (KIR), within-*N. wattersi* (WATT), and between species (BW) comparisons. Positive normalized enrichment scores (NES) indicate enrichment toward the high-FST end of the ranked gene list (divergent), whereas negative NES indicate enrichment toward the low-FST end (conserved). The table highlights major interpretable themes rather than providing an exhaustive or strictly rank-ordered list; all significant categories are reported in Table S2.

| Contrast | Direction | Functional theme | Representative GO term | NES | FDR | Representative lead-edge module |
| --- | --- | --- | --- | --- | --- | --- |
| KIR | Divergent | Translation initiation | BP: Translational initiation (GO:0006413) | 2.05 | 0.029 | EIF family (10 genes), *gtf2f1*, *trmt6* |
| KIR | Conserved | Neural maintenance | BP: Myelination (GO:0042552) | -2.08 | 7.79 x 10<sup>-4</sup> | *tuba8l3* |
| WATT | Divergent | Bioenergetics | BP: ATP biosynthetic process (GO:0006754) | 2.14 | 0.004 | ATPase family (7 genes) |
| WATT | Conserved | Oxygen transport | Oxygen transport (GO:0015671) | -2.31 | 2.04 x 10<sup>-5</sup> | *cygb*, *gb-α2*, *gb-β3*, *gb-β4*, *hbg1*, *ngb* |
| WATT | Conserved | Chromatin organization | Nucleosome assembly (GO:0006334) | -2.57 | 7.81 x 10<sup>-8</sup> | *h1m*, *hist1h1t*, *set* |
| WATT | Conserved | Transcription initiation | BP: DNA-templated transcription, initiation (GO:0006352) | -1.90 | 0.040 | *eif2aka1*, *gtf2f1*, *trmt6* |
| BW | Divergent | DNA binding | MF: Double-stranded DNA binding (GO:0003690) | 2.03 | 0.031 | *fos*, FOX family (8 genes), *gata1a*, *mterf*, *tp53* |
| BW | Divergent | Transcription factor binding | Transcription factor binding (GO:0008134) | 1.87 | 0.043 | *dip2a*, *dip2b*, FOX family (10 genes), *psmc5*, *snai1a* |

Leading-edge analyses identified subsets of genes contributing disproportionately to each enrichment signal (Fig. S2, Table S7). Relative to genome-wide averages, leading-edge genes exhibited elevated mean exon divergence for translation initiation in *N. kirki* (1.74x the genome-wide mean F_ST_, ATP biosynthesis in *N. wattersi* (1.60x), and double-stranded DNA binding and transcription factor binding in the interspecific comparison (1.56x and 1.59x, respectively). In contrast, genes associated with myelination (0.21x), nucleosome assembly (0.43x), and oxygen transport (0.41x) in *N. wattersi* exhibited substantially lower divergence than the genomic background (Fig. 3b). Leading-edge genes associated with significant enrichments were distributed across multiple chromosomes rather than concentrated within individual genomic regions (Fig. 4). In the interspecific comparison, enrichment was driven primarily by members of the FOX gene family together with *psmc5*, *snai1a* and other regulatory genes (Fig. 4, Table 2).

**Figure 4:**
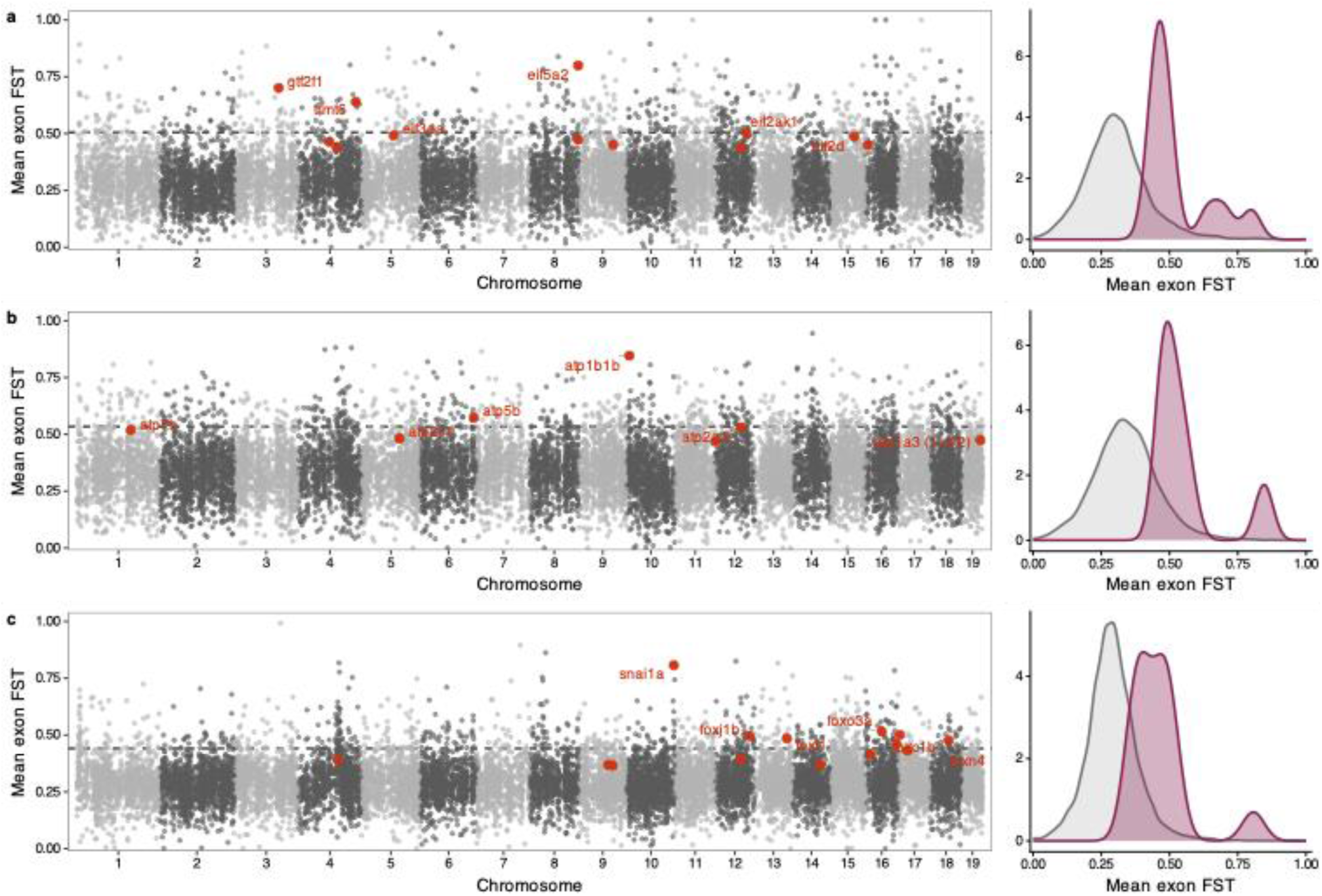
Genomic distribution of leading-edge genes underlying enriched functional themes. Gene-level exon differentiation (mean exon FST per gene) plotted across the 19 assembled chromosomes of *Nothobranchius furzeri*. Grey points represent all annotated protein-coding genes, with alternating shades indicating adjacent chromosomes. Red points indicate leading-edge genes contributing to the corresponding GSEA signal, and labels identify the most differentiated leading-edge genes: **(a)** translation initiation in *N. kirki*, **(b)** ATP biosynthetic process in *N. wattersi*, and **(c)** transcription factor binding in the interspecific comparison. Associated density plots show the distribution of mean exon FST values for leading-edge genes (purple) relative to the genome-wide background (grey).

Unsupervised phylogenetic reconstruction based on genome-wide SNP variation provided an independent assessment of genomic structure (Fig. 5). Without specifying species or population identity, individuals formed two strongly supported clades corresponding to *N. kirki* and *N. wattersi*. Within each species, the phylogeny further resolved well-supported geographic subclades that closely mirrored hydrological organization (Fig. S4). In *N. kirki*, the upland Namwera population formed a deeply divergent lineage sister to a Lake Chilwa-Lake Chiuta clade. Within *N. wattersi*, major clades broadly corresponded to the upper Shire Valley, the eastern and western shores of Lake Malawi, and northern Lake Malawi populations (Fig. S5), with additional substructure among neighboring localities. These phylogenetic relationships closely matched the PCA-based clustering (Fig. 2), independently recovering both species boundaries and geographically structured population divergence.

**Figure 5:**
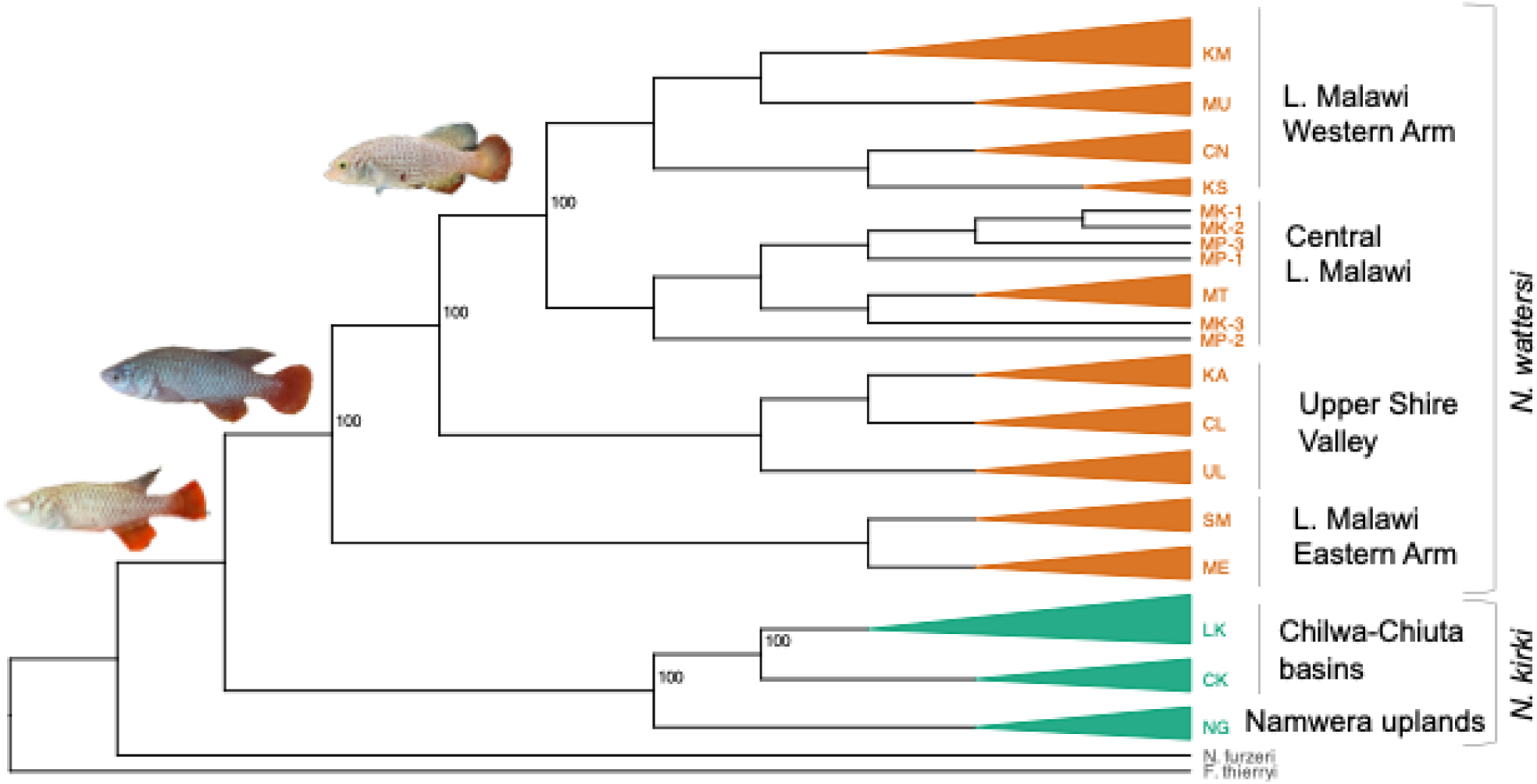
Population-level phylogeny inferred from 46 whole-genome sequence data using WASTER. Monophyletic populations were collapsed into triangles, with one representative label per population. Populations belonging to *N. kirki* are shown in green and those belonging to *N. wattersi* in orange. Individual samples are retained for populations that were not recovered as monophyletic (MK and MP). *Fundulopanchax thierryi* and *N. furzeri* were included as outgroups for rooting. Numbers at selected internal nodes indicate branch support values.

#### Morphological divergence

Morphological analyses revealed that variation was structured more strongly among populations within species than between species. Before morphometric analyses, length-mass allometric relationships were evaluated across species and populations. Significant log(*TL*) x species (*F*_1, 976_ = 7.32, *p* = 0.0069) and log(*TL*) x population (*F*_17, 944_ = 4.16, *p* = 3.47 x 10^−8^) interactions indicated heterogeneous allometric scaling relationships, supporting the use of population-aware size correction procedures. Following size correction, the combined PCA explained 70.0% and 10.1% of total variation along PC1 and PC2, respectively (Fig. S3). Species did not differ significantly along PC1 (β = 0, p = 1.0), and morphospace occupancy overlapped extensively between species, indicating that morphological differentiation was structured primarily among populations rather than between species.

The dominant axis of morphological variation represented coordinated changes across multiple anatomical regions rather than variation in a small subset of traits. In both species, PC1 showed relatively uniform contributions from head, body, fin, and positional measurements, indicating an integrated axis of body-form variation following size correction (Fig. S6, Fig. S7). Secondary axes captured more localized patterns of variation. In *N. kirki*, these involved fin-position and proportional traits, whereas in *N. wattersi*, PC2 was driven primarily by jaw and snout dimensions, indicating additional craniofacial differentiation largely independent of the primary body-form axis.

### Population structure and geographic organization

#### Genomic structure

At the population level, genome-wide SNP variation revealed additional spatial structure within both species. In *N. kirki*, individuals clustered according to geographic origin, separating populations associated with Chilwa, Chiuta, and the Namwera Uplands. In *N. wattersi*, genomic clusters broadly corresponded to drainage systems associated with the Lake Malawi floodplains and the Shire basin, with additional substructure among local Lake Malawi populations (Fig. 2). These patterns indicate that hydrological geography structures genomic variation within both species.

Genome-wide differentiation among populations was moderate for both species. Within *N. kirki*, the mean pairwise F_ST_ across loci was 0.181 (median = 0.088), whereas within *N. wattersi* (Fig. S8) the mean was slightly higher at 0.208 (median = 0.106). Window-based analyses again revealed heterogeneous differentiation: using the 99th percentile threshold, highly divergent windows were defined by F_ST_ > 0.335 in *N. kirki* and > 0.371 in *N. wattersi*, with 171 windows exceeding the high-divergence threshold in each species.

#### Morphological structure

Morphological variation likewise exhibited pronounced population structure within both species. PC1 - PC3 explained 60.3%, 9.6%, and 8.4% of morphological variance, respectively (78.3% cumulative). *N. kirki* populations from the Chiuta basin occupied strongly positive PC1 scores while those from the Chilwa basin clustered on the negative side of the axis (Fig. 6a). PERMANOVA confirmed a highly significant population effect (F_1,17_ = 19.05, R^2^ = 0.544, p = 0.001), while tests of multivariate dispersion were non-significant, indicating that population divergence reflected differences in multivariate means rather than unequal dispersion. In *N. wattersi*, populations occupied distinct regions of morphospace, with several populations separating primarily along PC1 and others exhibiting additional differentiation along PC2 (Fig. 6b). PERMANOVA confirmed strong population-level divergence (F_8,171_ = 20.40, R^2^ = 0.50, p < 0.001), and most pairwise population comparisons were significant. These results indicate that spatial organization, especially hydrological context, is a major axis of morphological divergence.

**Figure 6:**
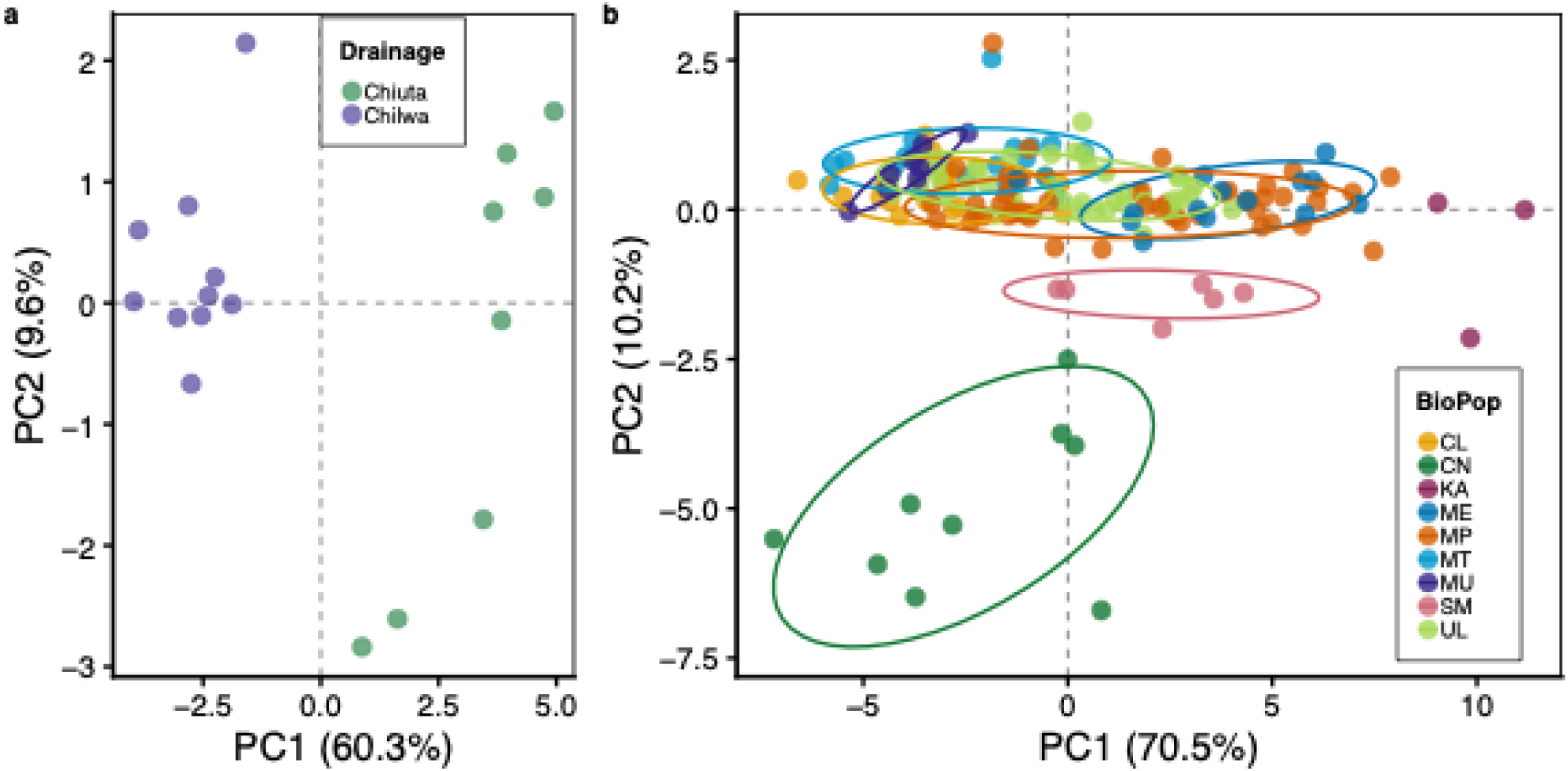
Principal components analysis of morphological traits: (**a**) Morphological variation in *N. kirki* from Lake Chilwa (green) and Lake Chiuta (orange). (**b**) Morphological variation in *N. wattersi* sampled across Lake Malawi and the Shire River floodplain.

#### Microbiome structure

Gut microbiome composition exhibited weaker but detectable spatial structuring. Alpha diversity (Observed richness and Shannon diversity) also varied among populations but showed substantial overlap, indicating considerable within-population heterogeneity (Fig. S9). Relative abundance patterns at the phylum level revealed population differences in microbial community composition, albeit with substantial within-population variation (Fig. 7). Principal coordinates analysis of beta diversity further showed partial clustering of individuals by species and population, broadly consistent with patterns observed in the genomic and morphological datasets. However, microbiome structure was less discrete and exhibited greater overlap among populations, suggesting that microbial communities reflect both host-associated structure and local environmental influences (Fig. S10a, Fig. S10b). These results indicate that microbiome composition is partially aligned with host population structure while remaining more labile than genomic or morphological divergence.

**Figure 7:**
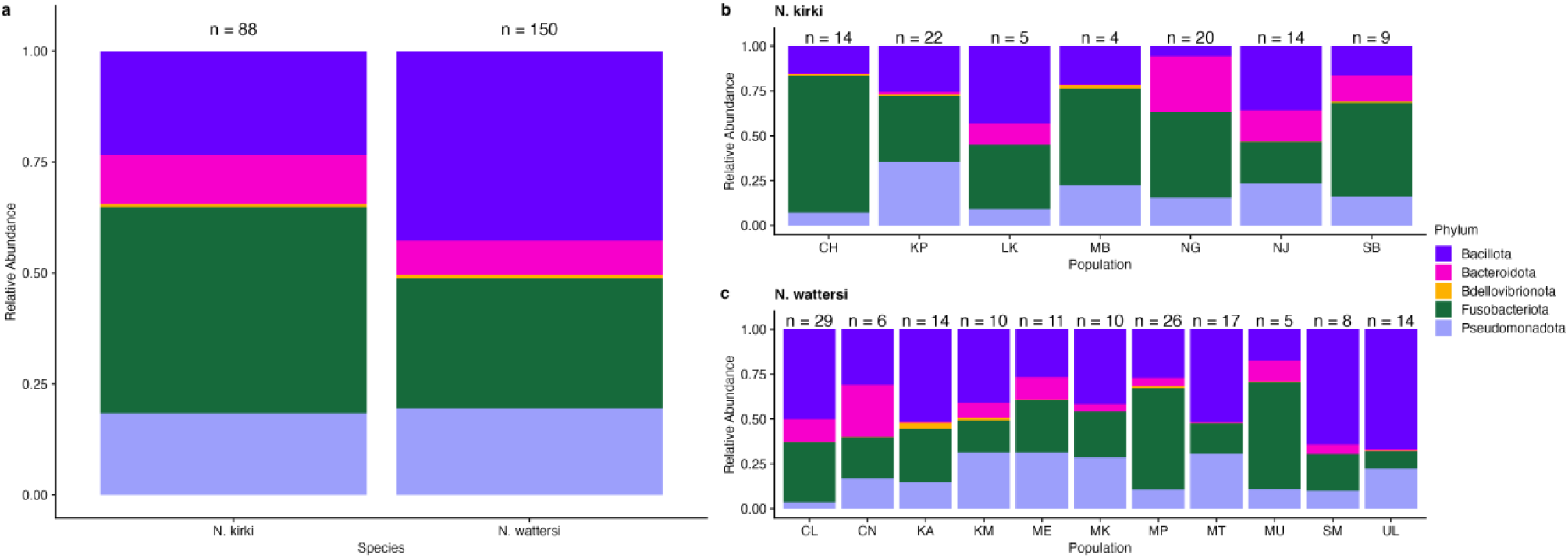
Relative abundance of the five most prevalent microbial phyla in gut samples. (a) Interspecific comparison between *N. kirki* and *N. wattersi*; (b) population-level variation within *N. kirki*; and (c) within *N. wattersi*. Across species, both taxa are dominated by similar major phyla but differ in their relative proportions, with *N. wattersi* showing a greater contribution of Bacillota and reduced representation of Fusobacteriota compared to *N. kirki*. Within species, microbiome composition is more variable among populations, with shifts in dominant phyla suggesting strong local environmental or host context effects, particularly evident in the heterogeneous profiles among *N. wattersi* populations. Note: Reduced representation of Bacillota coupled with increased Bacteriota in the upland Namwera (NG) *N. kirki* population.

### Functional architecture of genomic divergence

Gene-level analyses of divergent genomic regions suggested that functional signal is broadly distributed rather than concentrated within a single pathway. Across the full gene sets, hundreds of genes were associated with divergence within *N. kirki*, within *N. wattersi*, and between species, but no GO categories remained significant after correction for false discovery rate. After restricting attention to terms supported by at least three genes and reducing redundancy, recurring themes involved cellular structural organization, cytoskeletal assembly, developmental processes, and neural processes. These results suggest that gene-window divergence is functionally diffuse across many biological processes.

Exon-level analyses contrasted with this diffuse window-based signal by revealing a strong concentration of coding divergence in transcriptional regulators. The leading-edge subset driving GSEA enrichment comprised 514 genes (Table S8), and transcription factors were strongly overrepresented among the most divergent genes (5.8% of top-ranked genes vs. 0.66% in the genomic background; Fisher’s exact test odds ratio = 9.27, p < 2.2 x 10^−16^). Rather than being confined to a few genomic regions, these genes were distributed across multiple chromosomes, with modest clustering of several FOX family members but no evidence that the enrichment was driven by a small number of genomic hotspots (Fig. 4c). Divergence is disproportionately concentrated in transcription factor families, indicating that regulatory evolution is a major component of genomic differentiation.

Morphological divergence was likewise integrated across coordinated multivariate trait combinations rather than concentrated in a small number of diagnostic measurements. In both species, major axes of variation represented integrated covariance among head, body, fin, and positional traits, indicating that population differentiation involves broad shifts in overall body form rather than isolated trait changes. Similar to the genomic results, morphological differentiation was distributed across multiple trait combinations and hierarchical levels of organization, with substantial variation occurring among populations within species and comparatively limited separation at the species level. These patterns show that phenotypic divergence is expressed as integrated multivariate differentiation rather than discrete shifts in single morphological characters.

### Environmental correlates of morphological divergence

Despite substantial morphological differentiation at population level, neither geographic distance nor genome-wide genetic distance explained patterns of morphological divergence. In *N. wattersi*, Mantel tests did not support isolation by distance, and a direct comparison between genomic and morphological population distances across the subset of populations with overlapping data was also non-significant (Mantel r = −0.021, p = 0.47). Thus, populations that are more genomically divergent are not necessarily more morphologically divergent.

In contrast, environmental variables showed a significant association with morphological variation. In *N. wattersi*, redundancy analysis of population mean morphology against elevation, water temperature, and pH explained 65.3% of total variance in the constrained component, and the global model was significant (F = 3.77, p = 0.025). The first canonical axis accounted for 74.2% of environmentally explained variation, and elevation emerged as the only significant predictor (F = 7.08, p = 0.012), whereas temperature and pH were not significant. These results suggest that environmental variation, particularly elevational differences among sites, may contribute to morphological divergence at the population level. Among the environmental variables examined, elevation showed the strongest association with morphological variation in the unconstrained analyses.

To assess whether associations between morphology and environment were influenced by shared population history, we conducted population-level redundancy analyses and variance partitioning using environmental principal components and genomic principal components derived from whole-genome SNP data. Environmental effects on morphology were not significant either before (RDA: F = 1.15, p = 0.352, adjusted R^2^ = 0.033) or after accounting for genomic structure (partial RDA conditioned on genomic PC1: F = 1.19, p = 0.325, adjusted R^2^ = 0.044; genomic PC1-2: F = 1.07, p = 0.347, adjusted R^2^ = 0.021). Variance partitioning indicated only a small unique contribution of environmental predictors to morphological variation (adjusted R^2^ = 0.021), whereas genomic predictors explained little additional variation after accounting for environmental effects (adjusted R^2^ = −0.114). The shared contribution of environmental and genomic predictors was likewise small (adjusted R^2^ = 0.012). After accounting for shared genomic structure, neither environmental nor genomic predictors explained substantial unique variation in morphology, indicating that morphology-environment associations were weak at the population level.

### Sex-specific effects on morphology

Sexual dimorphism contributed significantly to multivariate morphological variation in *N. wattersi*. A PERMANOVA testing sex alone showed that sex accounted for 9.1% of total multivariate variance (R^2^ = 0.091, F = 17.08, p = 0.001). When sex and population were included additively, the model explained 69.9% of the total variation (R^2^ = 0.699, F_10_,_171_ = 37.41, p = 0.001), indicating that both factors contribute strongly to overall morphological variation. In contrast, the sex-by-population interaction was not significant (R^2^ = 0.017, F_6,171_ = 1.58, p = 0.099), providing little evidence that the magnitude or pattern of sexual dimorphism differs consistently among populations. Tests for multivariate homogeneity of dispersion showed no differences between sexes (permutational test, p = 0.135), indicating that sex effects primarily reflect shifts in multivariate trait means rather than differences in within-sex variability. Consistent with the trait distributions (Fig. S11), males and females exhibited parallel differences across populations, although the magnitude of dimorphism varied among individual traits. Sexual dimorphism therefore represents a robust and conserved component of morphological variation superimposed on population level differentiation.

**Figure 8:**
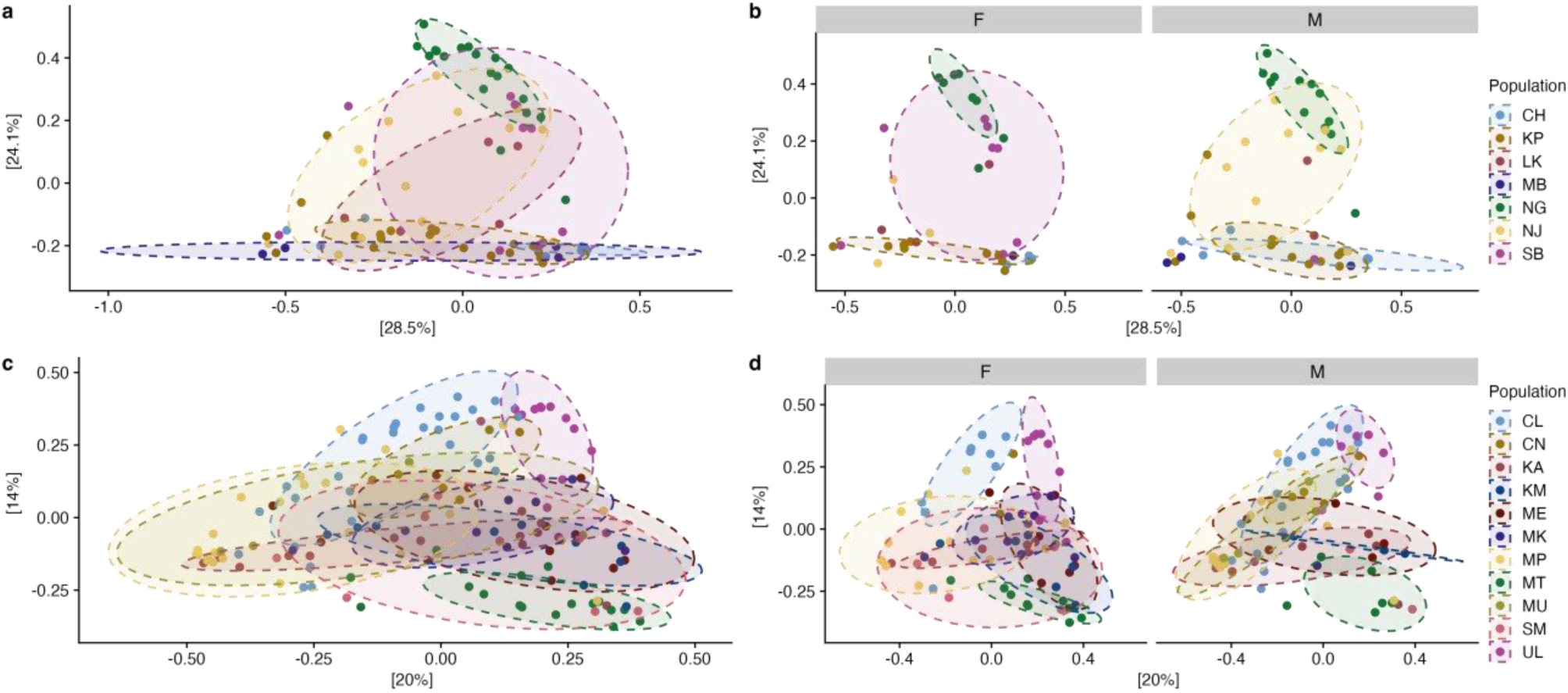
Beta diversity of gut microbial communities in wild fish based on Bray-Curtis distances. Principal coordinates analysis (PCoA) of gut microbiome composition for *N. kirki* (top row) and *N. wattersi* (bottom row). Left panels (**a**, **c**) show inter-population differences, while right panels (**b**, **d**) show samples stratified by sex (F = female, M = male). Each point represents an individual fish, colored by group (see legend). Ellipses denote 75% confidence regions around group centroids. Across both species (left panels), samples cluster by population, indicating structured differences in microbial community composition among groups. When stratified by sex (right panels), population-level clustering is largely retained, although differences in dispersion and orientation suggest potential sex-specific variation in microbiome structure.

## Discussion

In this study, we integrated data from whole genome sequencing, multivariate morphology, population-level phylogenetic structure, and gut microbiome composition across matched populations of *N. kirki* and *N. wattersi* to evaluate if divergence occurs with a consistent structure across biological levels. Consistent with earlier work identifying these taxa as distinct evolutionary lineages (Ng’oma et al., 2013), we show that geographic and hydrological structure strongly organize genomic differentiation within and between species. Furthermore, we found that genomic, phenotypic, and microbiome divergence are only partially coupled, indicating that different biological layers capture distinct components of evolutionary divergence. Notably, exon-level analyses revealed distinct functional signatures across evolutionary scales, including translation-associated processes within *N. kirki*, energetic and ion homeostasis functions within *N. wattersi*, and transcriptional regulatory functions in the interspecific comparison. Together, these results suggest that genomic divergence is not functionally uniform but instead involves different biological pathways at different hierarchical levels of divergence.

### Hierarchical divergence and partial coupling across biological levels

Divergence in *Nothobranchius* is hierarchically structured across biological levels, but coupling among these levels is incomplete and context dependent. Consistent with predictions of the G2E framework, genome-wide differentiation shows the clearest and most stable spatial structuring, strongly separating species and populations and reinforcing the phylogenetic structure recovered in PCA and FST analyses (see Fig. 5). In contrast, phenotypic and microbiome divergence are only partially aligned with this genomic framework. Morphological differentiation is substantial within species and broadly correlated with biogeographic distribution, yet morphological distances do not track genome-wide differentiation in a simple monotonic fashion, and neither geographic distance nor broad environmental gradients fully explain phenotypic variation (compare Fig. 6 with Figs. 2 & 5). Similarly, microbiome composition reflects aspects of host species and drainage structure but exhibits substantially greater overlap among populations within species than genomic clustering.

This weaker population structure is consistent with microbial communities integrating more recent ecological conditions than host genomic variation, while still retaining detectable signatures of host evolutionary history. Comparable patterns have been reported in other systems, where phenotypic divergence is only weakly associated with genome-wide differentiation despite strong population structure, e.g. (Leinonen et al., 2013), and where host-associated microbial communities respond more rapidly to local environmental conditions than host genomic structure (Foster et al., 2017; Henry et al., 2021). Together, these studies and our results suggest that genomic, phenotypic, and microbiome divergence may integrate ecological and evolutionary processes over different spatial and temporal scales.

Our sensitivity analyses indicate that accounting for population relatedness does not materially alter inference regarding morphological divergence. Although morphology varied substantially among individuals and populations, at the population level, morphology exhibited only modest and scale-dependent associations with environmental variation, and little correspondence with genome-wide genomic structure. Conditioning environmental effects on genomic relatedness produced nearly identical results, suggesting that the observed patterns are unlikely to reflect environmental responses confounded by shared evolutionary history.

These patterns suggest that different biological layers capture divergence through distinct but overlapping ecological and evolutionary processes. Genome-wide differentiation provides a stable spatial framework associated with long-term hydrological isolation and historical connectivity, whereas phenotypic variation reflects additional influences of sex, allometry, developmental constraints, ecology, and potentially behavioral variation. Microbiomes represent an even more environmentally responsive layer shaped by both host-associated and local ecological factors. This intermediate position between host genomic structure and local environmental conditions illustrates why microbiomes provide a complementary perspective on evolutionary divergence rather than a direct surrogate for host genetic differentiation. Consistent with this view, the dominant bacterial phyla observed in *N. kirki* and *N. wattersi* differed from those previously reported in wild *N. furzeri* populations, where Proteobacteria dominated (Smith et al., 2017), whereas Fusobacteria and Bacillota (Firmicutes) were often dominant in the populations examined here (Fig. 7). Although such comparisons should be interpreted cautiously because of differences among species, environments, and study designs, they further illustrate the dynamic nature of host-associated microbial communities. Taken together, our findings support a hierarchical model in which geographic and hydrological structure generate stable genomic differentiation, while phenotypic and microbiome variation represent increasingly context-dependent and only partially aligned layers of divergence (Fig. 9).

**Figure 9.**
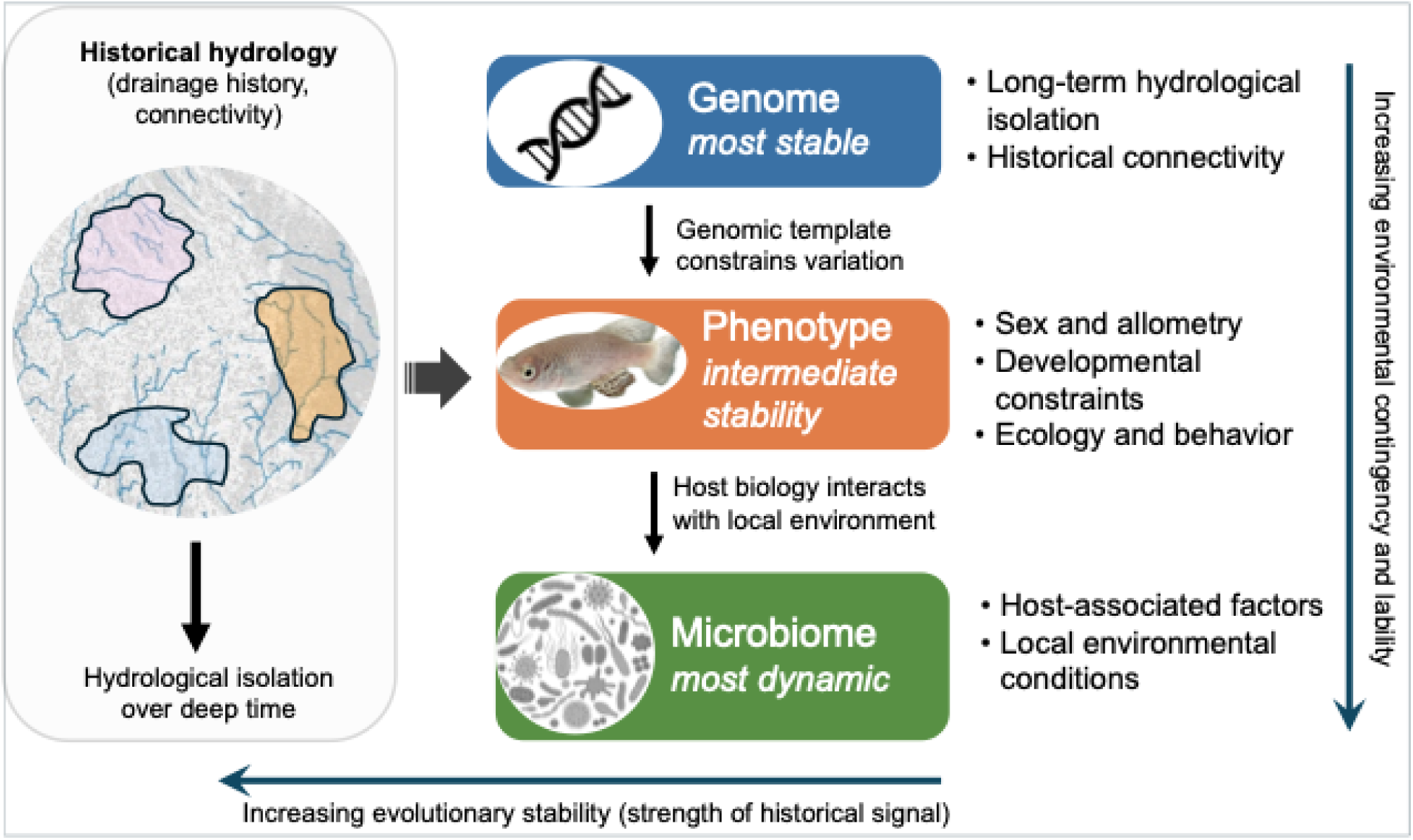
Conceptual model of hierarchical divergence across biological levels in hydrologically structured landscapes. Hydrological isolation establishes a stable genomic framework on which phenotypic and microbiome divergence accumulate through increasingly context-dependent processes.

### Hydrological structure as a backbone of divergence

Geographic and hydrological structure generate stable genomic differentiation that forms the primary spatial framework for divergence, while phenotypic and microbiome variation represent more context-dependent and only partially aligned biological layers shaped by ecology, sex, plasticity, and local environmental conditions. Addressing our first objective, the results support a hierarchical model of divergence in which hydrological configuration provides a geographic backbone for divergence across biological levels. Within this framework, genomic differentiation shows the clearest and most consistent spatial structuring, while phenotypic and microbiome variation are only partially aligned with this underlying pattern. Genome-wide variation clearly separates both species and populations, and this structure is independently supported by phylogenetic clustering, which recovers the same major divisions without prior information on species or population identity. Morphological analyses similarly reveal strong population-level differentiation, with basin- and drainage-associated shifts in multivariate trait space. Microbiome composition, while more variable, also shows partial alignment with structure at species and population levels. Taken together, these results indicate that geographic configuration and hydrological connectivity are central organizing axes of divergence across multiple biological levels.

Between-species divergence is most plausibly linked to the geological history of the Lake Malawi Rift basin. Subsidence of the basin beginning in the late Miocene (∼8.6 Ma), together with uplift of the eastern rift shoulder, reorganized regional drainage and isolated the Chilwa-Chiuta systems from the Lake Malawi-Shire basin (Lancaster, 1981; Shroder, 1972; Watters, B.R., 1991c). The divergence between *N. kirki* and *N. wattersi* likely reflects this prolonged tectonic and hydrological isolation associated with Neogene to early Pleistocene restructuring of the East African Rift system. In contrast, divergence within *N. kirki* between Chilwa and Chiuta populations is expected to be relatively recent. The two lakes were connected prior to the Pleistocene and likely remained intermittently connected until as recently as ∼44 ka (Lancaster, 1981; Thomas et al., 2009). Today, they are separated by a relatively narrow east-west sand ridge (∼30 km long, 100 m - 1 km wide), which includes gaps near its western end where streams may have flowed into the late 19th century (Nicholson, 1998). Such a recent and potentially permeable barrier suggests the possibility of ongoing or historically recent gene flow through episodic hydrological connectivity or alternative dispersal mechanisms.

A notable example of finer scale within-species spatial structuring is the strong divergence of the upland Namwera population from lowland Chilwa-Chiuta populations within *N. kirki*. Although geographically proximate to the Chiuta system, this population occupies a distinct topographic setting and exhibits substantial genomic differentiation, suggesting that local hydrological and elevational barriers may generate additional hierarchical structure beyond major basin divisions. Similar geographically localized structuring is also evident within *N. wattersi*, where genomic clustering broadly follows drainage configuration and regional topography.

### Functional signatures of divergence across evolutionary scales

Functional enrichment analyses revealed that different evolutionary scales were associated with distinct biological processes. Within *N. kirki*, differentiated genes were enriched for translation initiation and protein synthesis functions, whereas differentiation within *N. wattersi* was associated primarily with ATP biosynthesis, ion transport, and oxygen transport pathways. Such functional heterogeneity is increasingly recognized across diverse taxa, where evolutionary change proceeds through different physiological and cellular mechanisms depending on environmental context and evolutionary timescale. For example, genomic studies of fishes and other vertebrates frequently report enrichment of metabolic, energetic, and ion-transport functions during adaptation to environmental gradients, particularly those involving temperature, oxygen availability, and salinity (Berg et al., 2015; Moreira & Smith, 2023; Wang et al., 2022). The coexistence of multiple functional axes of differentiation within *N. wattersi* is likewise consistent with patterns found in other systems, further supporting the working hypothesis that adaptation often involves concurrent changes across several biological pathways rather than a single dominant mechanism (Maharjan et al., 2012). These results suggest that population divergence may proceed through a mosaic of functionally distinct pathways, providing the physiological flexibility necessary to respond to heterogeneous and changing environments. Notably, nucleosome assembly exhibited strong negative enrichment within *N. wattersi*, indicating conservation of core chromatin-packaging functions despite differentiation in energetic and ion-transport pathways. This pattern is consistent with divergence being concentrated in specific physiological processes while fundamental genome organization functions remain under strong evolutionary constraint.

In contrast, the strongest interspecific enrichments involved transcription factor activity and transcription factor binding, suggesting that divergence between species is concentrated disproportionately in genes occupying regulatory positions within cellular networks. This pattern is consistent with a growing body of evidence indicating that evolutionary divergence frequently involves modifications to regulatory architecture and gene expression networks rather than changes restricted to downstream structural genes (Signor & Nuzhdin, 2018; Stern & Orgogozo, 2008; Uller et al., 2018). Regulatory divergence has been implicated in phenotypic evolution across diverse taxa and may facilitate coordinated changes in multiple traits through effects on shared developmental and cellular pathways (Holloway et al., 2007; Jacobs et al., 2024; Mack & Nachman, 2017).

Particularly notable was the recovery of multiple FOX family transcription factors among the most differentiated leading-edge genes, including *FOXO3A, FOXJ1B, FOXL1, FOXC1B,* and *FOXN4*, together with the zinc-finger transcriptional regulator *SNAI1A*. FOX genes regulate diverse developmental, physiological, and homeostatic processes across vertebrates (Hannenhalli & Kaestner, 2009), whereas *SNAI1* occupies a central position in cellular differentiation and developmental signaling networks (Dong & Wu, 2021). Although direct genotype-phenotype links cannot be established from the present data, the repeated recovery of transcription-associated functions and regulatory genes suggests that evolutionary changes in regulatory architecture may have contributed to species divergence, potentially influencing multiple downstream developmental, physiological, and metabolic pathways. Importantly, leading-edge regulatory genes were distributed across multiple chromosomes rather than concentrated in a small number of genomic regions, supporting a polygenic architecture of divergence. These observations support the hypothesis that divergence between species is associated with coordinated changes in regulatory architecture rather than isolated changes in individual structural genes.

### Implications for multi-level evolutionary inference

Our results show that divergence in *Nothobranchius* is hierarchically structured across genomic, phenotypic, and ecological dimensions, but that coupling among these layers is incomplete and context dependent. Genome-wide differentiation captures historical isolation and long-term spatial structure, whereas phenotypic and microbiome variation reflect additional developmental and environmental influences operating over different ecological contexts. These findings emphasize that no single biological layer fully captures the complexity of evolutionary divergence. Instead, integrating genomic, phenotypic, and ecological data provides a more complete understanding of how environmental heterogeneity shapes diversification across biological scales.

Two limitations remain important. First, direct genotype-phenotype relationships are not resolved, and future work linking regulatory divergence to specific trait combinations will be essential.

Second, incomplete sample overlap among genomic, morphological, and microbiome datasets limits direct matrix-based comparisons across all populations. Additional sampling, finer-scale environmental measurements, and integration of behavioral and life-history data would help clarify the processes linking divergence across levels. Expanding microbiome analyses beyond broad taxonomic summaries to functional or strain-level resolution may also reveal stronger links to host divergence.

## Acknowledgments

Authors thank Manuel Leal, Kevin Middleton, Elizabeth King, and members of the King and Ng’oma laboratories for constructive criticism on the manuscript.

## Data Accessibility and Benefit-Sharing

### Data accessibility Statement

Raw whole-genome and 16S rRNA amplicon sequencing reads, and associated sample metadata are deposited in the NCBI Sequence Read Archive (SRA) under BioProject ID PRJNA1490097. Morphological trait data are provided with this manuscript (Tables S1a and S1b).

### Benefit-Sharing Statement

This research was conducted in compliance with the Nagoya Protocol on Access and Benefit Sharing. All research and material transfer permits were obtained from the Government of Malawi. The study contributes knowledge relevant to the conservation and evolutionary biology of Malawi’s native annual killifishes and freshwater biodiversity. It forms part of a long-term research collaboration led by a Malawian scientist in partnership with Mzuzu University (MZUNI) and other Malawian collaborators. As part of this collaboration, fieldwork provided hands-on training in annual killifish ecology for Malawian students and research staff. In addition, project support enabled a week-long symposium on annual killifish biology, ecology, and conservation for MZUNI graduate students and faculty in Malawi. The authors are committed to equitable international scientific partnerships, research training, and institutional capacity building.

## Author Contributions

EN conceived the study and collected samples. TRJ performed whole-genome sequencing and morphological data collection. PKW performed 16S rRNA amplicon sequencing and microbiome analyses. AWT conducted phylogenetic inference from whole-genome sequencing data. EN performed the genomic and morphological analyses. EN wrote the manuscript. All authors contributed to manuscript revision and approved the final version. The study was supported by a University of Missouri start-up award to EN.

## Ethics approval

Fish used in this study were collected in the field under permits issued by the Government of Malawi. All procedures involving animals were conducted in accordance with the guidelines of the University of Missouri Institutional Animal Care and Use Committee (IACUC) under approved protocol #63571.

