## Supplemental Figures for "Hierarchical divergence across genomic, phenotypic, and microbiome dimensions in two annual killifish species from Malawi"

**Table of Contents:**

| Figure S1. SNP density vs gene-level divergence | Page 2 |
| --- | --- |
| Figure S2. **Divergence distributions of leading-edge genes** | Page 3 |
| Figure S3. PCA of morphometric traits | Page 4 |
| Figure S4. Whole-genome-inferred phylogeny | Page 5 |
| Figure S5. Phylogeny annotated PCA of genomic variation | Page 7 |
| Figure S6. **Trait loadings of the combined morphometric PCA** | Page 8 |
| Figure S7. Biplots of morphological traits driving divergence | Page 9 |
| Figure S8. **Distribution of gene-level exon divergence** | Page 10 |
| Figure S9. Alpha diversity | Page 11 |
| Figure S10. Phylum-level relative abundance in **a.** *N. kirki*, **b.** *N. wattersi* | Page 12 |
| Figure S11. Alpha diversity by sex | Page 13 |

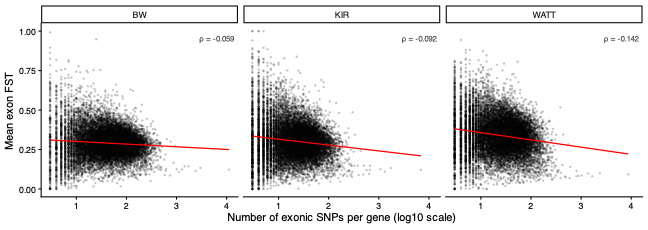

**Figure S1. Relationship between exon SNP density and gene-level divergence.** Mean exon FST per gene plotted against the number of exonic SNPs assigned to each gene for the between-species (BW), within-N. kirki (KIR), and within-N. wattersi (WATT) comparisons. Red lines indicate fitted linear trends. Gene-level divergence exhibited only weak negative correlations with exonic SNP count (Spearman ρ = - 0.059 to - 0.142), indicating that variation in SNP density explained little of the observed patterns of exon-level differentiation.

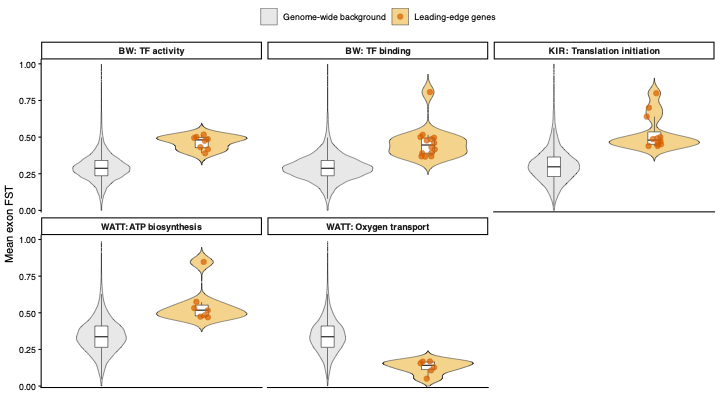

**Figure S2. Divergence distributions of leading-edge genes relative to genome-wide background genes.** Violin plots show the distribution of mean exon FST values for all genes included in each contrast (grey) and leading-edge genes contributing to significant GSEA enrichments (gold). Red points indicate individual leading-edge genes. Panels correspond to transcription factor activity and transcription factor binding in the between-species comparison (BW), translation initiation in N. kirki (KIR), and ATP biosynthesis and oxygen transport in N. wattersi (WATT).

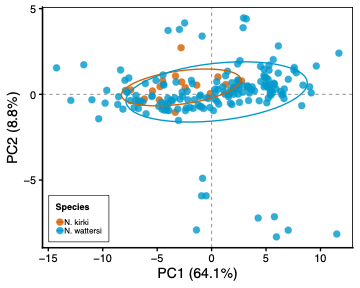

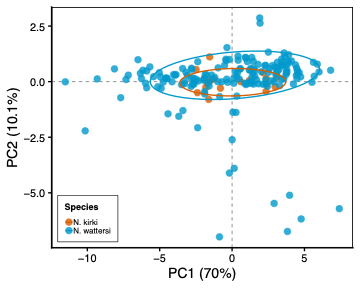

**Figure S3.** Principal components analysis (PCA) of a combined size-corrected morphometric traits from *N. kirki* (n = 18) and *N. watersi* (n = 172). Species occupied largely overlapping regions of multivariate morphospace, indicating limited morphological differentiation at the species level relative to within-species variation.

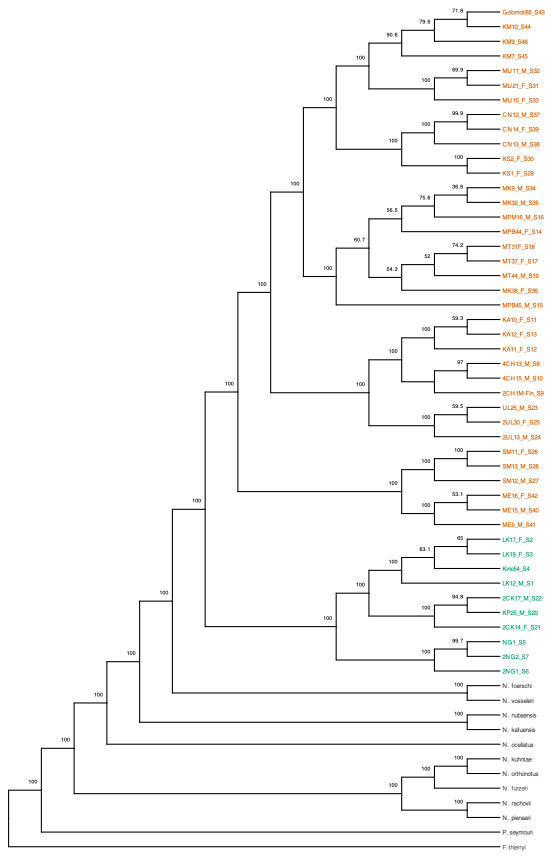

**Figure S4.** Complete whole-genome phylogeny inferred using WASTER, including all 58 samples analyzed. The tree includes all reference taxa, 46 individual samples from all populations, and branch support values for all internal nodes. Fundulopanchax thierryi was used to root the tree. Population labels are color-coded by species (N. kirki, green; N. wattersi, orange), whereas reference taxa are shown in black.

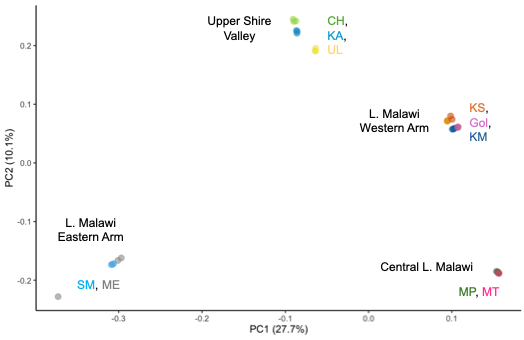

**Figure S5.** Phylogeny annotated principal components analysis (PCA) of genomic variation within *N. wattersi*. Populations cluster geographically by drainage, identifying four separate clusters: 1) Shire Basin, 2) West L. Malawi Arm, 3) East L. Malawi Arm, and 4) Northern Group (Salima).

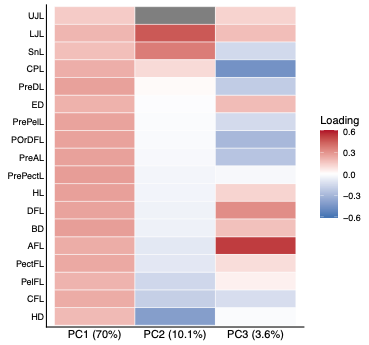

**Figure S6. Trait loadings for the first three axes of the combined *N. kirki* and *N. wattersi* morphometric PCA.** PC1 received positive contributions from all size-corrected traits, indicating that most morphological variation was aligned along a common axis of variation in body form. PC2 and PC3 captured secondary variation in cranial, fin, and body proportions.

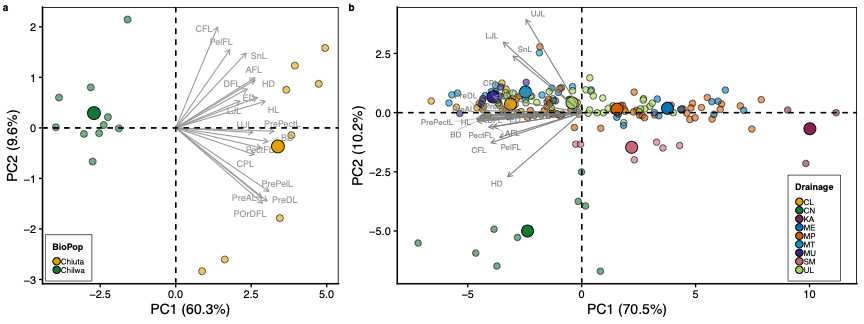

**Figure S7.** Biplots of morphological variables superimposed on PCA plots showing morphological traits driving divergence among populations within each species: **a.** *N. kirki* L. Chiuta and L. Chilwa basins; and **b.** *N. wattersi* in L. Malawi and Shire River flood plains. Small points are individual samples; large points are population centroids showing relative population separation in multivariate morphospace.

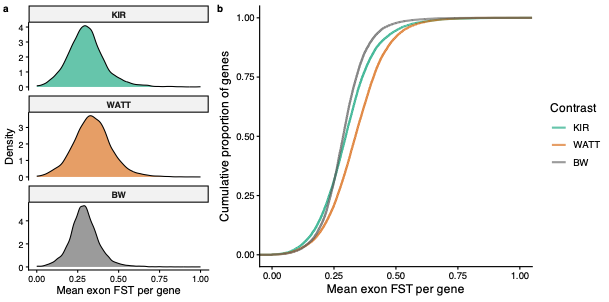

**Figure S8. Distribution of gene-level exon divergence across evolutionary scales.** (a) Density distributions and (b) empirical cumulative distribution functions (ECDFs) of mean exon FST values across all annotated protein-coding genes retained for exon-level analyses. Mean exon FST was calculated from all exonic SNPs assigned to each gene. Curves are shown separately for within-N. kirki (KIR), within-N. wattersi (WATT), and between-species (BW) comparisons and illustrate the underlying gene-level divergence distributions used for gene set enrichment analyses (GSEA).

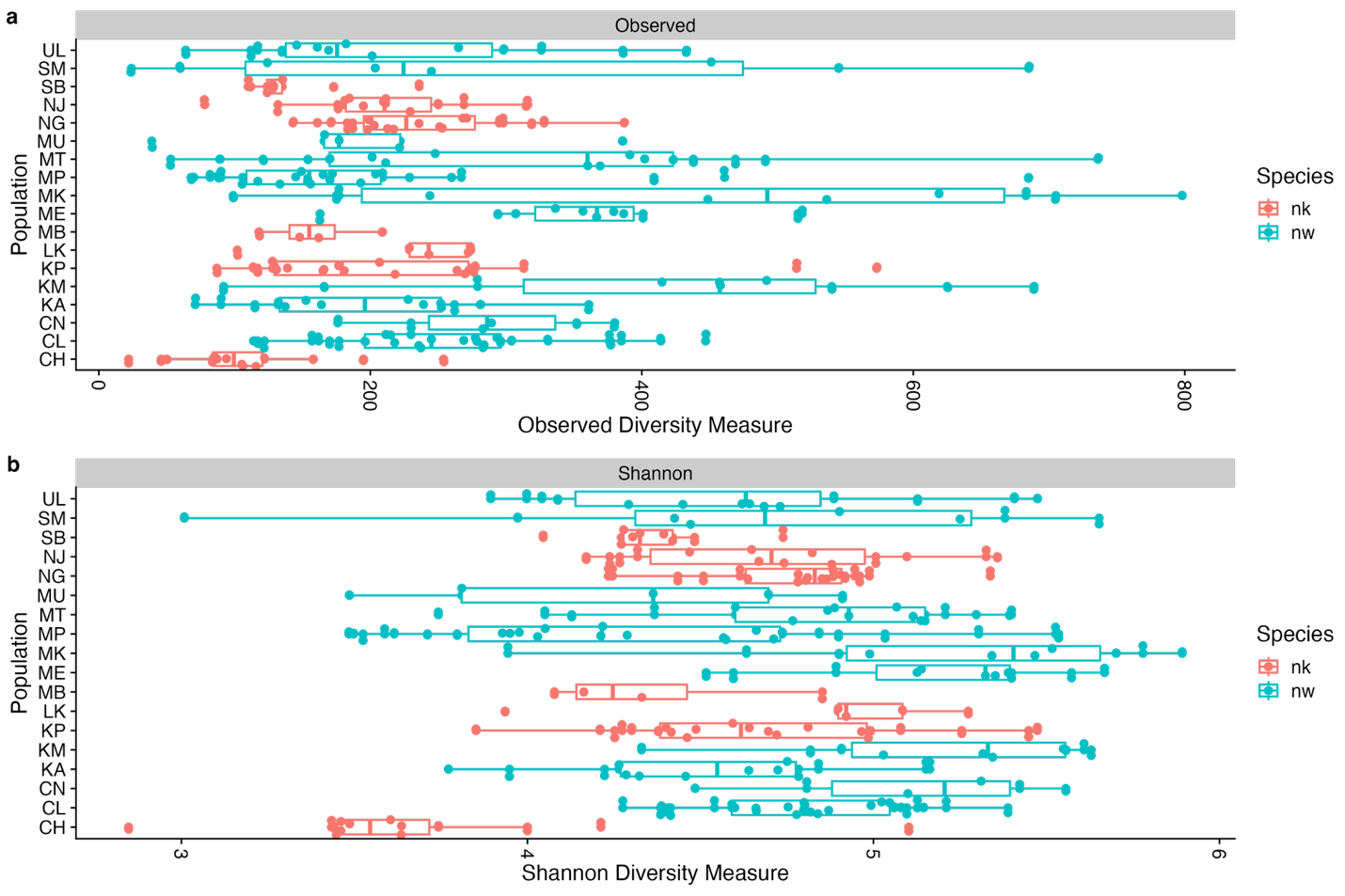

**Figure S9. Alpha diversity of gut microbiomes across populations of** N. kirki **(nk) and** N. wattersi (nw)**.** Observed richness (top) and Shannon diversity (bottom) are shown for each population. Boxplots indicate the median and interquartile range, whiskers extend to 1.5 x the interquartile range, and points represent individual fish. Populations are grouped by species.

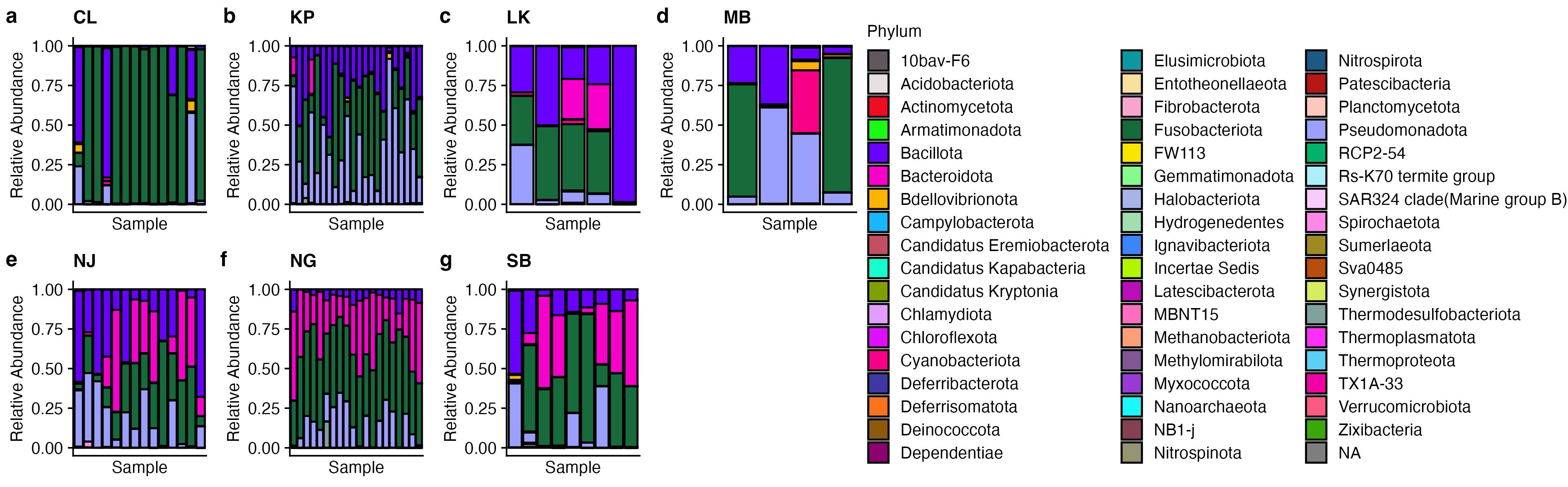

**Figure S9.** Phylum-level relative abundance log scale of bacterial communities across populations of *N. kirki*. Each bar represents the mean relative abundance of samples collected from that population. Colors correspond to bacterial phyla shown in the shared legend.

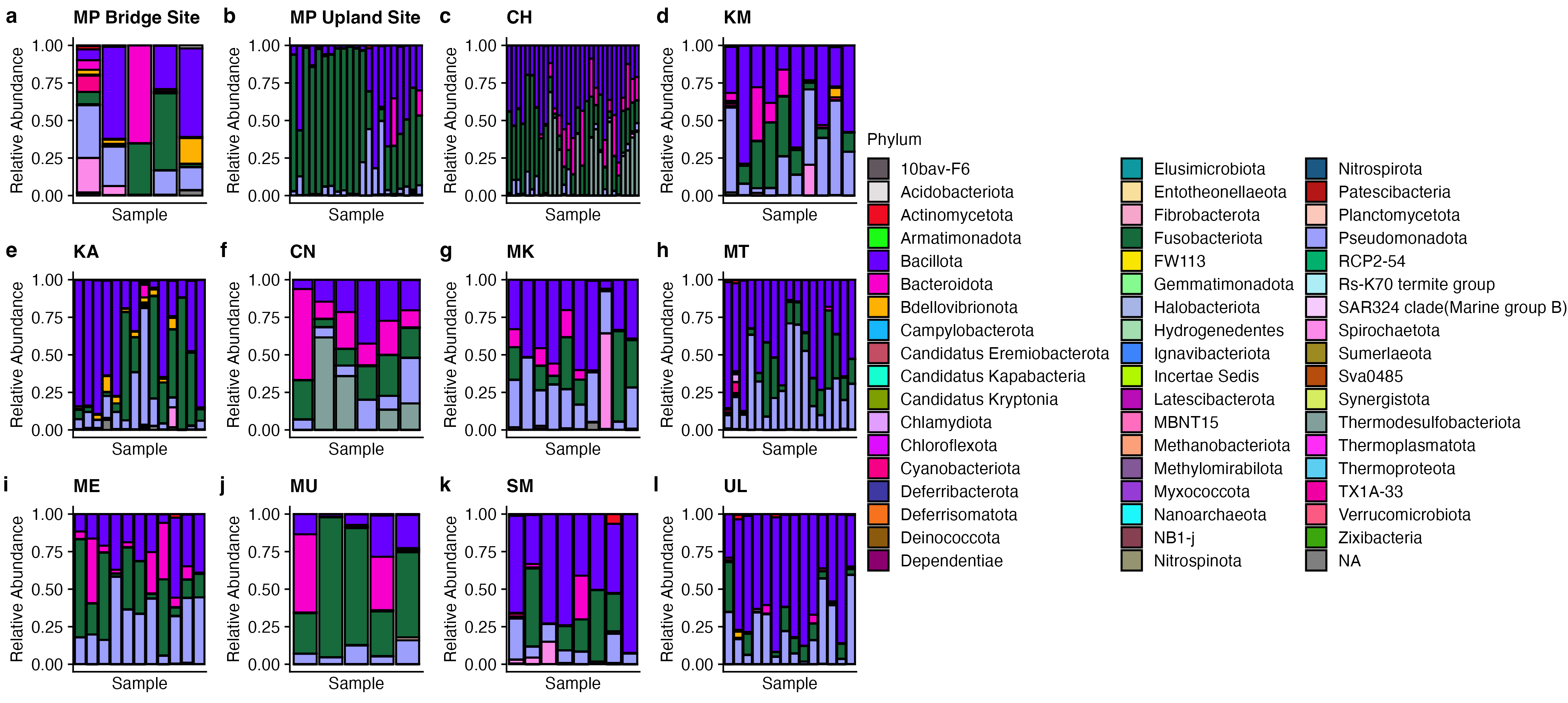

Figure S10: Phylum-level relative abundance log scale) of bacterial communities across populations of *N. wattersi*. Each bar represents the mean relative abundance of samples collected from that population. Colors correspond to bacterial phyla shown in the shared legend.

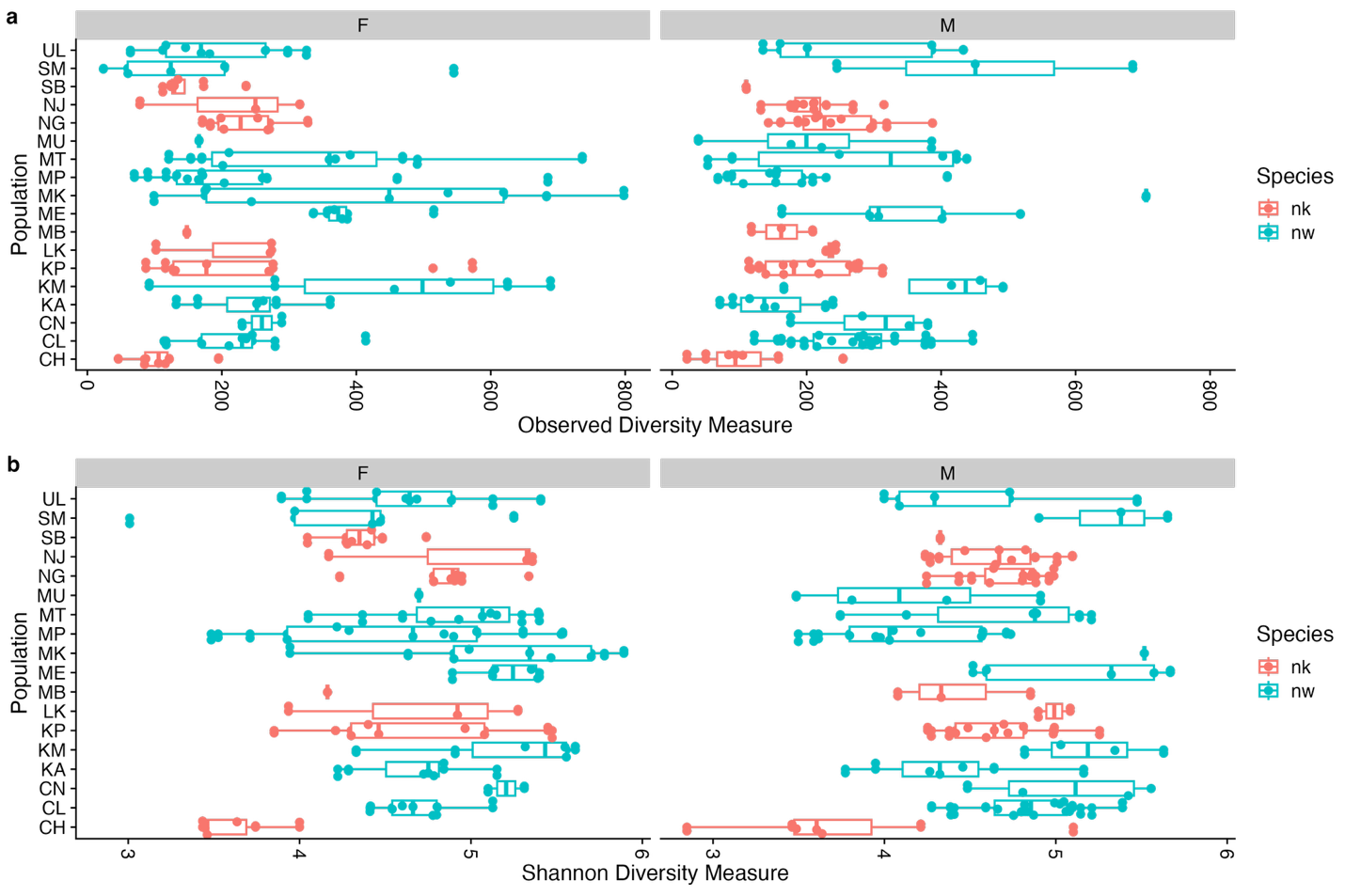

**Figure 11.** **Sex-specific patterns of gut microbiome alpha diversity across populations of** N. kirki **(nk) and** N. wattersi (nw)**.** Observed richness (top) and Shannon diversity (bottom) are shown separately for females (F) and males (M). Boxplots indicate the median and interquartile range, whiskers extend to 1.5 x the interquartile range, and points represent individual fish.
